# Life history traits predict the contribution of transient dynamics to variation in population growth

**DOI:** 10.64898/2026.08.28.747639

**Authors:** Hung-wei Lin, Christina M. Hernández, Harman Jaggi, Wenyun Zuo, Shripad Tuljapurkar, Roberto Salguero-Gómez

**Author notes:** **Corresponding mailing address**: Department of Biology, Life and Mind Building, University of Oxford, OX1 3EL, Oxford, United Kingdom.

## Abstract

The performance of any natural population in variable environments depends on contemporaneous changes in its vital rates (*e.g.*, survival, reproduction) as well as legacies carried by its population structure. Yet whether the relative contribution of these two pathways can be predicted from life history remains unknown. Here, we use stochastic simulations of 1,986 matrix population models from 137 species to quantify the contribution of transient dynamics to variation in population growth rate, and test its associations with key life history traits. Longer generation times were associated with reductions in transient contributions, contrary to theoretical expectations. Greater stage-specific survival heterogeneities were associated with increases in transient contributions, whereas greater iteroparity was associated with decreases in plants but increases in animals. These associations were robust to body size, phylogenetic relationships, and vital-rate variability. Life history traits therefore provide a strong predictor for when population structure shapes population responses to environmental variability.

## Introduction

Understanding what drives variation in population growth is central to both ecological inference and management (Sibly et al., 2005). This fundamental question is intimately linked to the increasing interest in resilience in ecology, since the resilience of natural populations can be reflected in how their growth rates respond to environmentally-induced changes in vital rates (*e.g.,* survival, reproduction) (Koons et al., 2009; Neubert & Caswell, 1997). A substantial body of literature has therefore examined how environmental variation influences vital rates (Ehrlén et al., 2016; Morris et al., 2008), how variation in these rates determines population growth rate (Caswell, 2001; Ellner & Rees, 2006), and how life history traits shape these links (Gascoigne et al., 2025; Koons et al., 2009, 2016; Salguero-Gómez et al., 2016a). However, to date studies have mostly considered environmental variation as temporary deviations from an otherwise constant baseline, either explicitly (*e.g.,* Capdevila et al., 2020) or implicitly (Crone et al., 2011). Consequently, we are yet to fully capture how persistent or recurrent disturbances generate variation in population growth through multiple pathways, including both the immediate effects of current disturbances on vital rates and the accumulated effects of past disturbances (Koons et al., 2016; McDonald et al., 2016).

Environmental fluctuations in natural systems are the rule rather than the exception. Under such fluctuating conditions, the structure of a population (*i.e.,* the distribution of individuals among state classes such as age or stage; Caswell, 2001) may fail to converge to a stable state distribution (Tuljapurkar, 1984). These persistent deviations from a stable population structure becomes particularly important under ongoing global change, which alters disturbance regimes (Turner, 2010; Wolkovich et al., 2014), intensifies environmental variability (Boer et al., 2000; Räisänen, 2002), and reduces the likelihood that populations experience prolonged periods near demographic equilibrium (Koons et al., 2005; Williams et al., 2011). Accordingly, analyses of variation in population growth rate increasingly account for deviations between transient (short-term) and stable population structure (long-term) rates (Capdevila et al., 2020; Jaggi et al., 2026; Morozov et al., 2024).

Fluctuations in realised population growth rate in stochastic environments arise from two intrinsic pathways in structured populations (Tuljapurkar, 1990). The first is the immediate pathway driven by time-varying vital rates. The second pathway is the historical pathway driven by departures of population structure from its stable stationary distribution accumulated from past variation in vital rates — referred to as ‘transient growth rate’ (Koons et al., 2005). Previous work in population ecology has primarily examined how environmental change alters contemporaneous vital rates and thereby influences population growth (Crone et al., 2011; Dunham et al., 2021; Jenouvrier et al., 2012). Yet variation in population growth rates is explained by both contemporaneous vital-rate changes as well as from transient growth driven by population structure (McDonald et al., 2016; Tuljapurkar, 1990). A large contribution of transient dynamics implies that realised growth rates are strongly influenced by structural legacies of past disturbances (Hernández et al., 2026). From a population recovery perspective, a high contribution of transient dynamics highlights the importance of the current structure of a population for its own growth. For instance, populations with too few reproductive individuals may recover slowly even when survival improves (White et al., 2013). Consequently, changes in environmental conditions that alter vital rates may not directly shape realised growth.

Previous studies have quantified the influence of transient dynamics on population growth from several angles (Ellis & Crone, 2013; Festa-Bianchet et al., 2003; Jiang et al., 2022; McDonald et al., 2016; Tuljapurkar et al., 2009). Measures such as damping time capture how quickly populations converge to their stable stationary structure, and thus the extent to which past perturbations influence current dynamics (Grant & Benton, 1996; Tuljapurkar, 1982). Comparative work has further shown that these measures are associated with life history strategies, with slower life histories exhibiting longer-lasting structural transient effects (Jiang et al., 2022; Tuljapurkar et al., 2009). Furthermore, using simulations, McDonald et al. (2016) attributed up to half of the stochastic dynamics of plant populations to transient dynamics. However, these authors measured the importance of transient dynamics with reactivity metrics (Neubert & Caswell, 1997) and correlation-based attribution. A key limitation is that this approach does not allow direct comparison of the contributions to population growth rate by population structure vs. contemporaneous variation in vital rates.

Recently, Hernández et al. (2026) developed an exact decomposition of realised population growth rate that separates the effects of immediate changes in vital rates from transient dynamics. Analytical results of their framework predict greater contributions to realised population growth rate from transient dynamics than from contemporaneous vital-rate fluctuations in populations with longer generation times and slower damping of population structure. However, whether this prediction holds under the observed magnitude of vital-rate variation across wild populations remains unknown. To address this gap, the decomposition framework and simulation-based approach of Hernández et al. (2026) allow us to empirically test how life history shapes the contribution of transient dynamics to population growth.

Here, we test hypotheses regarding how life history traits may predict the proportion of temporal variation in population growth rate arising from transient dynamics rather than from contemporaneous vital-rate fluctuations (hereafter, the ‘transient contribution’). Using 1,986 matrix population models (Caswell 2001) across 137 species from the COMPADRE and COMADRE databases (Salguero-Gómez et al., 2015, 2016b), we quantify the transient contribution and its association with three key life history traits, generation time, degree of iteroparity, and life-cycle stage specific survival heterogeneity (hereafter ‘survival unevenness’), while controlling for organismal size and phylogenetic ancestry. Motivated by life history theory, we predict that (H1a) longer generation time reduces temporal variation in population growth rate but (H1b) raises the transient contribution to such variation, because slow turnover in slow life histories dampens variation in growth and causes transients decay more slowly (Gaillard & Yoccoz, 2003; Jiang et al., 2022). We also predict that (H2a) greater degree of iteroparity lowers both temporal variation in growth rate and (H2b) the transient contribution, since narrow reproductive windows generate cohort pulses that amplify growth fluctuations and transient contributions (Botsford et al., 2014; Smallegange & Berg, 2019). Finally, we predict that (H3a) greater survival unevenness raises both temporal variation in growth rate and (H3b) the transient contribution, because departures from the stable stage distribution matter more when survival differs strongly among stages (Caswell, 2001, p. 246). Figure 1 summarises our predictions.

**Figure 1.**
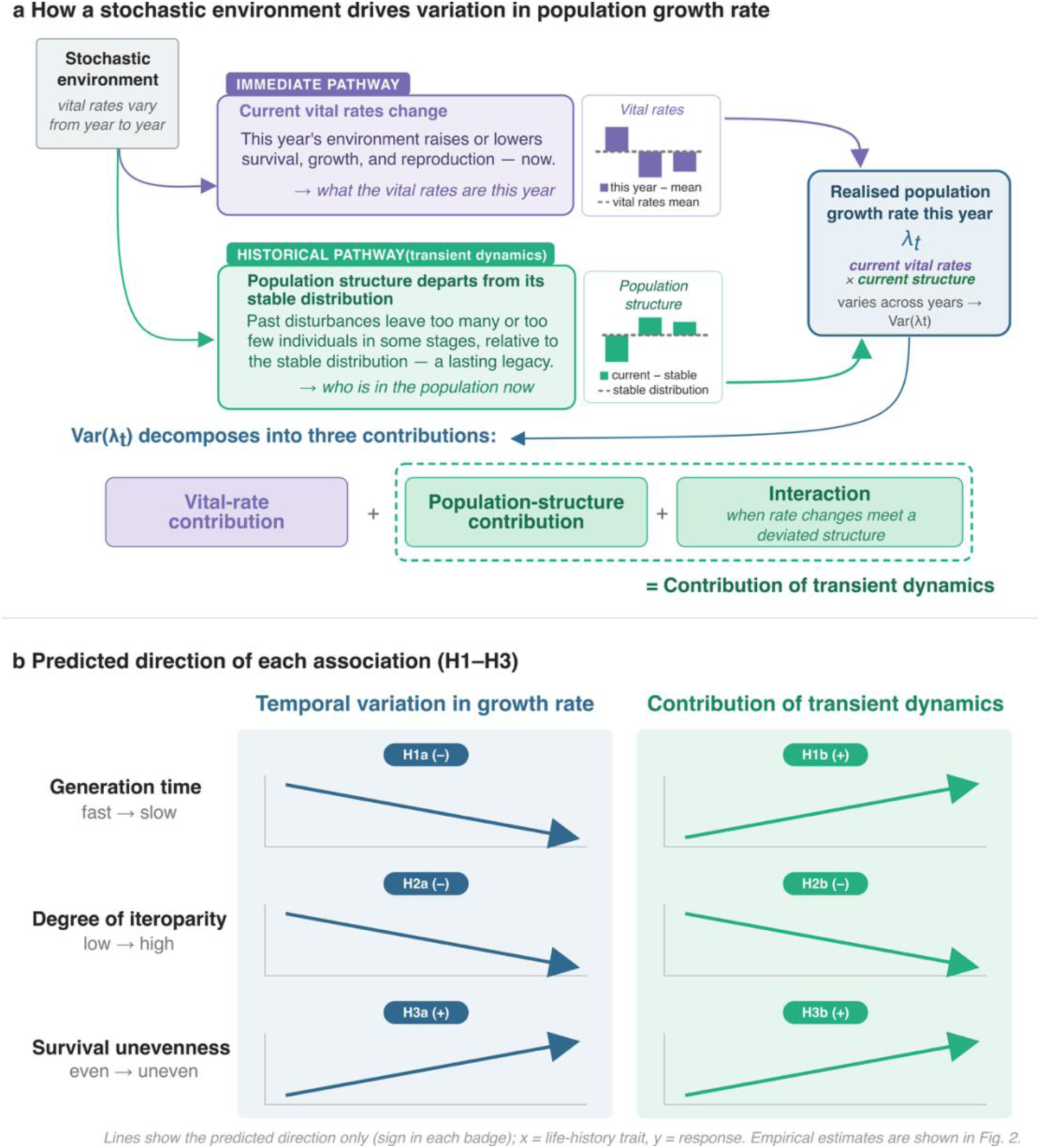
Conceptual framework linking a stochastic environment to variation in population growth rate, and the predicted role of life history traits. (a) In a stochastic environment, vital rates (survival, growth, reproduction) fluctuate from year to year, and this variation can shape the realised population growth rate in year *t* (*λ_t_*) through two non-mutually exclusive pathways. The immediate pathway (purple) is the direct effect of the current year’s vital rates, which deviate from their long-run mean (inset: this year vs. mean). The historical pathway (green) acts through the population structure: past disturbances can trigger the current stage distribution of the population to depart from its projected stable distribution — the fixed proportional distribution of individuals among stages that a population would settle into, and thereafter maintain, if vital rates remained constant (*i.e.,* the stable stage distribution; Caswell 2001). This departure is the legacy of past variation and is the source of transient dynamics. Realised population growth is obtained by considering how the current vital rates shape the current stage structure (current vital rates × current structure), so its temporal variance, Var(*λ_t_*), can be decomposed into three contributions: a vital-rate contribution, a population-structure contribution, and higher-order interaction terms, such as arise when fluctuating vital rates act on a structure that already departs from the stable distribution. The two components involving population structure (dashed outline) are grouped as the contribution of transient dynamics. (b) Predicted associations between three life history traits — generation time (*T*), degree of iteroparity (*S*) and survival unevenness (*Vσ*) — and the two response variables: temporal variation in growth rate (Var(*λ_t_*); blue) and the proportional contribution of transient dynamics (green). Survival unevenness here is defined as the coefficient of variation in survival probability among life stages, so that higher values indicate greater disparity in survival across the life cycle. Arrows indicate the predicted direction of each association under hypotheses H1a–H3b (sign in parentheses); they convey direction only and are not fitted relationships.

## Methods

### Data Collection

To calculate variation in population growth rate in stochastic environments and life history traits, we used matrix population models (MPMs hereafter; Caswell, 2001). We obtained MPMs from the COMPADRE Plant Matrix Database version 6.23.5.0 (Salguero-Gómez et al., 2015) and COMADRE Animal Matrix Database version 4.23.3.1 (Salguero-Gómez et al., 2016b). These two databases contain demographic data compiled as age-, size-and/or developmental stage-structured MPMs.

To simulate the stochastic population dynamics, we used temporally replicated MPMs within each examined population. To that end, we filtered the databases to retain only natural, unmanipulated populations with at least three MPMs (*i.e.*, 4 time points) available from different time points for the same population. To ensure that transient dynamics could emerge, we further restricted analyses to MPMs with at least three life stages (Stott et al., 2011). Finally, we required that each MPM could be decomposed into survival-transition and fertility submatrices, allowing calculation of the life history traits in this study. After imposing these selection criteria and excluding five populations whose life history traits contained non-finite values, our resulting dataset comprised 1,986 MPMs from 343 populations, including 315 populations from 112 plant species and 28 populations from 25 animal species.

To ensure that differences in demographic variability reflected life history rather than body size, we included adult body size as a covariate, since larger-bodied taxa consistently exhibit lower demographic variability (Cohen et al., 2012). We used maximum height for plants and adult body mass for animals; sources for all species are detailed in Table S1 and the compilation procedure in Supplementary Methods.

### Variation in population growth rate

To quantify temporal variation in population growth rate, we analysed simulated population trajectories. We define variation in population growth rate as the temporal variance of the realised population growth rate (*λ_t_*) across the stochastic simulations.

We calculate the realised population growth rate for time step *t* as

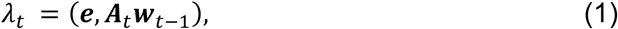

where ***A****_t_* is the population projection matrix at time *t*, ***w****_t_*_−1_ is the population structure at the start of the time step, and ***e*** is a vector of ones so that (***e***, ***A****_t_**w**_t_*_−1_) provides the total population growth across all stages.

We define the mean projection matrix as

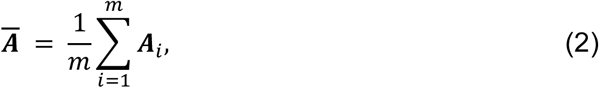

where *m* is the number of observed annual projection matrices for the population of interest. We then calculated the asymptotic growth rate *λ_d_* as the dominant eigenvalue of the mean projection matrix ***A̅***.

### Decomposing the variation in population growth rate

Following Hernández et al. (2026), we decompose the variation in population growth rate, we expanded the realised population growth rate *λ_t_* as

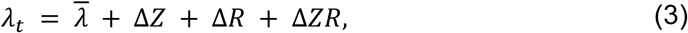

where *ΔZ* is the contribution to population growth rate arising from deviations in population structure, *ΔR* is the contribution arising from fluctuations in vital rates, and *ΔZR* is the interaction term.

Expanding realised population growth rate allows us to partition the variance in population growth rate into contributions from population structure, vital rates, and their interaction. The full variance partition is given by

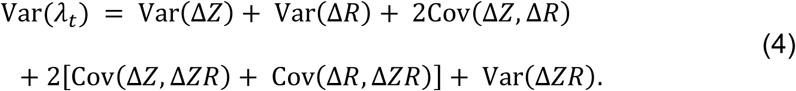

Following this decomposition, all terms except Var(*ΔR*) involve fluctuations in population structure. These terms therefore capture the effects of transient dynamics arising from deviations in population structure and their interaction with variation in vital rates. For clarity, we group these components (*i.e*., Var(Δ*Z*) + 2Cov(Δ*Z*, Δ*R*) + 2[Cov(Δ*Z*, Δ*ZR*) + Cov(Δ*R*, Δ*ZR*)] + Var(Δ*ZR*)) and refer to them collectively as the variation from transient dynamics (*V*_transient_). Thus, we can re-write the variance in population growth rate from equation 4 as:

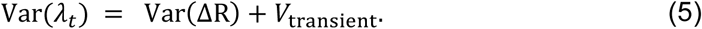

In a supplementary analysis we separately modelled the two components of the transient contribution, its structural component (Var(*ΔZ*)) and its interaction component, in the same way as our main analysis; we found that removing all terms other than structural component did not change our findings (Fig. S1–S2).

Finally, we defined the transient contribution (*C*_transient_) as the proportion of the total variance in realised population growth rate attributable to transient dynamics:

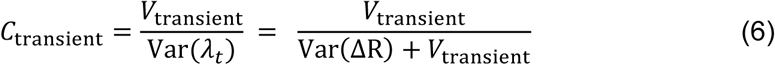

so that the vital-rate and transient contributions sum to one.

### Simulation of the stochastic environment

To quantify variation in population growth rate and transient contribution in stochastic environments, we used a framework developed by Hernández et al. (2026). Specifically, we simulated stochastic environmental variation in vital rates using the set of annual transition matrices available for each population (*e.g.*, repeated observations from the same site across multiple years). For each simulation, we generated a sequence of environments by randomly sampling from these matrices with equal probability, such that the observed temporal variation in vital rates defines the stochastic process. To estimate the transient contributions, we ran 500 independent simulation replicates per population, with a burn-in period to avoid the influence of the initial population structure (Hernández et al., 2026). Treating the decomposition terms as random variables across replicates, we estimated their variances and covariances to obtain the variance decomposition above (Equations 3–6); full detail of the simulations is given in Supplementary Methods.

### Life history traits

To test hypotheses H1a–H3b, we quantified three key life history traits for each population. These traits are generation time, degree of iteroparity, and unevenness in survival across stages from the mean projection matrix ***A***. We calculated generation time (*T*) as the average age difference between parents and offspring (Bienvenu & Legendre, 2015). Populations with low (high) *T* values are referred to as fast (slow) species (*sensu* Stearns, 1998). We calculated degree of iteroparity using Demetrius’ entropy (*S*) (1978), whereby larger values of *S* indicate longer reproductive windows. To calculate unevenness in survival across the life cycle (*V*_σ_), we first calculated stage-specific survival probabilities (*σ_j_*) as the probability that an individual in each life stage *j* survives to the next time step. We then calculated the coefficient of variation (CV) among these stage-specific survival values. Higher values of *V*_σ_ therefore indicate greater heterogeneity in survival probabilities along the life cycle in the examined population. The formulae for computing these three life-history traits are provided in the Supplementary Methods.

### Temporal variation in vital rates

To further investigate whether the associations between life history traits and the transient contribution (H1b–H3b) instead reflect differences in vital-rate variation, we quantified vital-rate variation for each population. To quantify such variation, we first identified the vital rate that most strongly influences the growth rate for a population, using the sensitivities of the mean projection matrix ***A*** (Caswell, 2001). We then calculated the CV of this rate across the population’s annual matrices; whereby higher values indicate larger year-to-year fluctuations in the most influential vital rate. We added this metric as a covariate and used the same phylogenetic multilevel model structure as the main analyses in a supplementary model.

### Phylogenetic corrections

To account for phylogenetic non-independence among species (Freckleton et al., 2002), we constructed plant and animal phylogenies and included their covariance matrices as a species-level random effect in our models. Tree construction and branch-length imputation are detailed in Supplementary Methods. Phylogenetic signal in both response variables, estimated as an analogue of Pagel’s *λ* (Pagel, 1999), was negligible in plants but higher and highly uncertain in animals (Fig. S3). We therefore report phylogenetically corrected models in the main text and uncorrected models in the Supporting Information (Fig. S4–S5).

### Statistical analyses

To test how life history traits shape the magnitude and source of variation in population growth rates (H1a–H3b), we fitted multivariate Bayesian multilevel models using the *brms* package (Bürkner, 2021) in R v4.4.2 (R Core Team, 2020). To optimise sampling efficiency and numerical stability, we ran each model for 8,000 iterations with a 50% warmup, and used conservative sampler settings to ensure reliable convergence (target acceptance rate = 0.975; maximum tree depth = 20; Bürkner, 2021).

To account for the correlation structure between the two response variables, we jointly modelled the variance in population growth rate (Var(*λ_t_*)) and the transient contribution. Because Var(*λ_t_*) was strongly right-skewed, we log-transformed this response prior to analysis. For all regression coefficients (*β*), we report posterior means and 95% credible intervals.

Predictors in each model comprised life history traits of interest alongside key confounders and random effects. For both plants and animals, the fixed effects included generation time *T*, degree of iteroparity *S*, and survival unevenness *V*_σ_. We additionally included two key factors known to affect these variables: log-transformed body size (plant height or animal body mass) and matrix dimension, as both can influence demographic variability independently of life history strategy (Salguero-Gómez et al., 2016a; Tenhumberg et al., 2009). To account for repeated species-level structure and variation in sampling effort (*i.e.*, the number of temporally replicated MPMs contributing to a population), we also included species identity and sampling effort as random intercepts. To reduce skew, we log-transformed generation time and survival unevenness, adding a small constant to the latter to accommodate zeros. To facilitate comparison of effect sizes among traits, we standardised all continuous predictors prior to modelling (Schielzeth, 2010).

To improve robustness to extreme values in the response variables, we fitted models with Student-t response distributions (Lange et al., 1989), and weakly informative priors throughout (Lemoine, 2019). Collinearity among predictors was low (all variance inflation factors < 3.1; Table S2; Zuur et al., 2010), and trait effects proved robust to leave-one-out sensitivity analyses in which we excluded each focal life history trait in turn. Priors, collinearity and sensitivity analyses (Fig. S6) are detailed in Supplementary Methods.

## Results

Life history predicts both temporal variation in population growth rate and the transient contribution to that temporal variation. Our models explain much of the variation in both responses: the conditional R^2^ is 0.66, 95% CI [0.60, 0.71] for the temporal variation in population growth rate and 0.57 [0.49, 0.63] for the transient contribution in plants, and 0.82 [0.61, 0.95] and 0.58 [0.38, 0.70] in animals, respectively. Fixed effects (*e.g.*, generation time, survival unevenness) alone explained a substantial share of both responses (marginal R^2^ = 0.25–0.53; Table S3). The fixed-effect signal is driven overwhelmingly by life history. The only covariate effects we detect are in plants, where larger body size and larger matrix dimension are weakly associated with greater temporal variation in population growth rate (*β* = 0.52, [0.04, 1.01] and *β* = 0.47, [0.06, 0.87], respectively; Table S3).

### Longer generation times reduce the transient contribution to variation in population growth rate

We find only partial support for our hypotheses H1, regarding the role of generation time in shaping both the magnitude of variation in population growth rate and the transient contribution to that variation. Consistent with our hypothesis H1a, which predicted that populations with longer generation time would exhibit less temporal variation in growth rate, we find this pattern across both taxa (plants: *β* = −1.76, 95% CI [−2.26, −1.23]; animals: *β* = −1.45, [−2.53, −0.30]; Fig. 2a, 2d). However, contrary to our hypothesis H1b, which predicted that longer generation time would raise the transient contribution to variation in growth rate, our results show that the transient contribution declines rather than increases with generation time in both plants (*β* = −0.06, [−0.11, 0.001]; Fig. 2a) and animals (*β* = −0.08, [−0.16, −0.002]; Fig. 2d).

**Figure 2.**
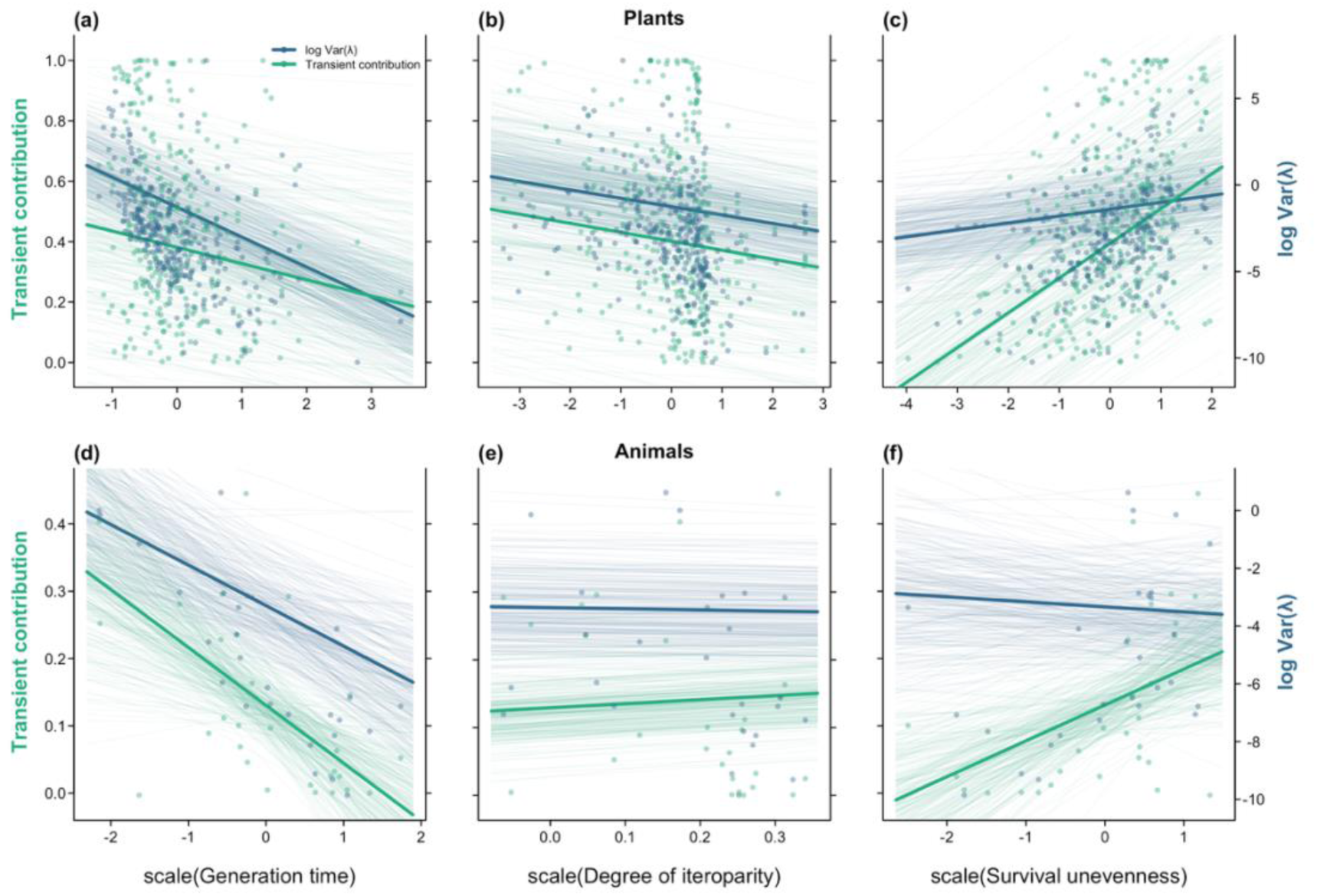
Life history predicts both the magnitude of variation in population growth rate and the contribution of transient dynamics to it: in plants, both decrease with generation time and iteroparity and increase with survival unevenness, whereas associations in animals are less certain and, for iteroparity, reversed. Each panel relates one life history trait (x-axis) to the proportional contribution of transient dynamics to the temporal variation in population growth rate (transient contribution; left y-axis, green) and log-transformed temporal variation in population growth rate (log Var(λ); right y-axis, blue). Because the transient contribution includes covariances between structural deviations and vital rate fluctuations, it can be negative; negative values indicate that population structure dampens, rather than amplifies, the variation in population growth rate generated by fluctuating vital rates. Panels show generation time (a, d), degree of iteroparity (b, e) and survival unevenness (c, f) for plants (a to c) and animals (d to f). All three life history traits were standardised to zero mean and unit variance before analysis, and the x-axes are on this standardised scale. Each point represents one population. Thin lines show 250 draws from the posterior of the full phylogenetic multilevel model, and thick lines the posterior mean. For visual clarity, three populations with extreme standardised trait values (|z| > 5; one each in panels a, b and e) are omitted from the plotted points and axis ranges; these populations were retained in all analyses, and the fitted lines shown reflect the full data. Axis tick labels appear only on the outer panels (transient contribution on a and d; log Var(λ) on c and f).

### Less iteroparous plants show a larger transient contribution to variation in population growth rate

We find support for our hypothesis H2 in plants but not in animals, regarding the role of iteroparity in reducing temporal variation in population growth rate and the transient contribution to that variation. As we had predicted via H2a and H2b (Fig. 1), we find that plant populations with higher degree of iteroparity exhibit both less temporal variation in growth rate (*β* = −0.48, 95% CI [−0.70, −0.26]; Fig. 2b) and a smaller transient contribution (*β* = −0.03, [−0.06, −0.003]; Fig. 2b). However, in our examined animal species, neither prediction is supported. Indeed, the association with temporal variation in growth rate is uncertain (*β* = −0.43, [−1.27, 0.40]; Fig. 2e), and the transient contribution increases rather than decreases with iteroparity in animals (*β* = 0.06, [0.01, 0.11]; Fig. 2e).

### More uneven survival across life stages raises the transient contribution to variation in population growth rate

We find support for our hypothesis H3 in plant and partial support in animals, regarding the role of survival unevenness on temporal variation in population growth rate and the transient contribution. As predicted via H3a and H3b (Fig. 1), plant populations with more uneven survival show greater temporal variation in growth rate (*β* = 0.39, 95% CI [0.10, 0.69]; Fig. 2c) and a larger transient contribution (*β* = 0.12, [0.08, 0.15]; Fig. 2c). In animals, only the transient contribution follows the predicted direction (Fig. 1b vs. Fig. 2f), with a credible interval that barely overlaps zero (*β* = 0.05, [−0.004, 0.11]; Fig. 2f), a notable result given the small sample. The association between survival unevenness and temporal variation in population growth rate remains uninformative (*β* = −0.14, [−0.92, 0.71]).

### Vital-rate variation does not account for the associations between life history traits and the transient contribution

To confirm that these trait associations do not merely reflect differences in vital-rate variability, in a supplementary model we added the temporal CV of the vital rate to which population growth rate is most sensitive as an additional covariate. That covariate shows no clear association with either response in plants or animals, and the associations of generation time, iteroparity, and survival unevenness with both responses remain qualitatively consistent with the primary analysis (Fig. S7, Table S4).

### Life history explains the weak covariation between temporal variation in growth rate and the transient contribution

The magnitude of temporal variation in population growth rate and the transient contribution to it are weakly related, and life history explains what they share. Across plant populations they covary weakly and positively before life history is accounted for (ρ = 0.13, 95% CI [−0.02, 0.28]; Fig. 3), yet this covariation disappears once predictors enter the model (ρ = 0.05, [−0.10, 0.20]; Fig. 3), because the same traits move both responses in tandem: survival unevenness raises both, while longer generation time lowers both. In animals the two responses are, if anything, negatively related, though too uncertainly to interpret (ρ = −0.17, [−0.67, 0.42] before and ρ = −0.28, [−0.71, 0.25] after accounting for life history; Fig. 3). Within-species variation in both responses was present but varied across species (Fig. S8).

**Figure 3.**
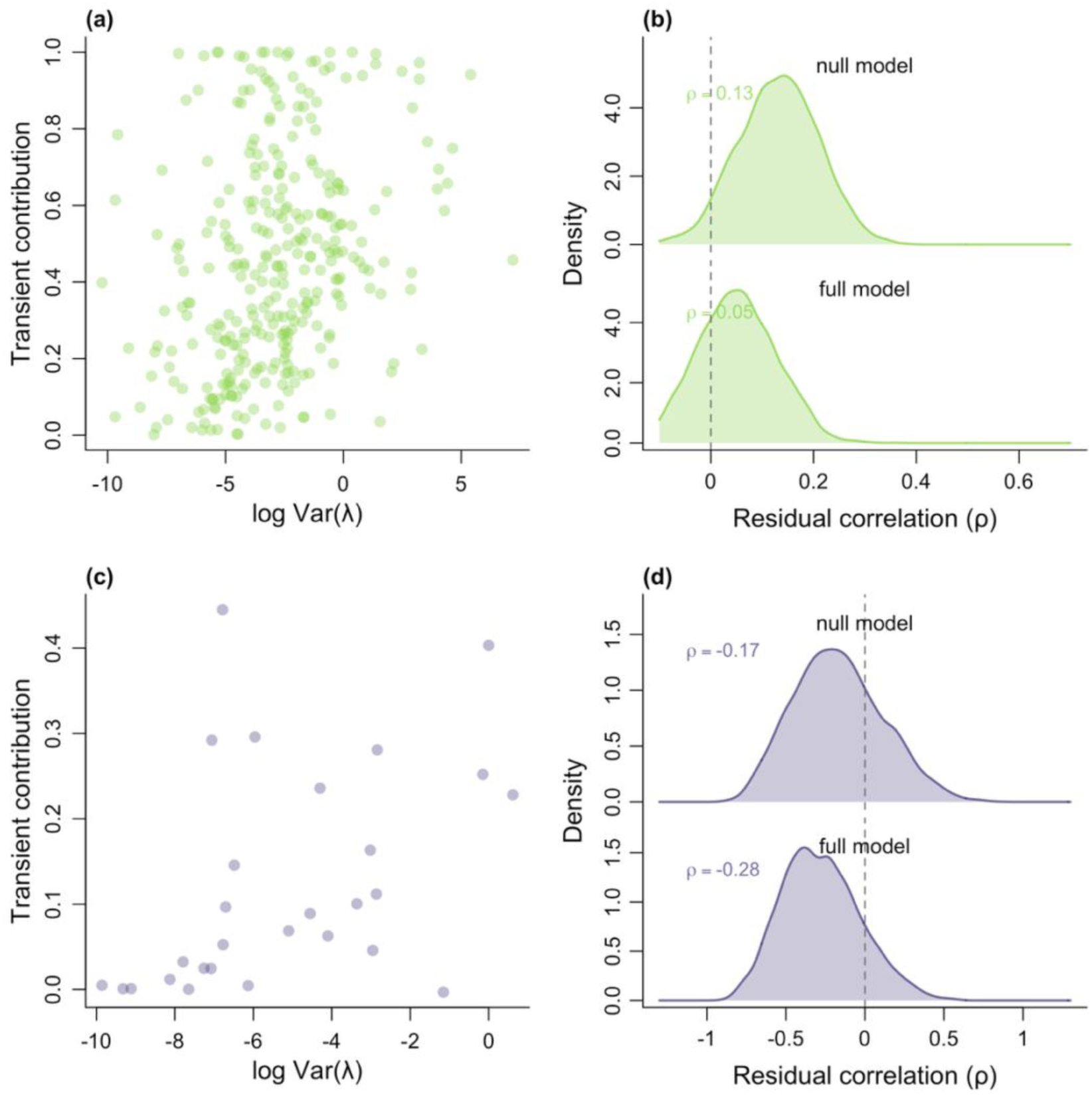
Plants show positive residual covariation between temporal variation in population growth rate and the transient contribution to that variation, and this covariation can be explained by life history, while animals show an uncertain negative association. Scatter plots of log-transformed temporal variance in population growth rate (log Var(λ); x-axis) against the proportional contribution of transient dynamics to that variation (transient contribution; y-axis) for plants (a) and animals (c); each point represents one population. As in Fig. 2, negative values of the transient contribution indicate that population structure dampens rather than amplifies variation in growth rate. Panels (b) and (d) show the posterior density of the residual correlation (ρ) between the two response variables from phylogenetic multilevel models, for plants and animals respectively. Reported ρ values are posterior means, and range from −1 to 1. The null model (intercept only; top) and the full model (including generation time, degree of iteroparity, survival unevenness, body size, and matrix dimension; bottom) are shown on a shared scale within each panel. The dashed vertical line indicates ρ = 0. After accounting for phylogenetic structure, plants show a weakly positive residual correlation under the null model (ρ = 0.13) that shrinks to near zero under the full model (ρ = 0.05), whereas animals show a negative but uncertain residual correlation (null: ρ = −0.17; full: ρ = −0.28), with considerable posterior uncertainty given the small animal sample size.

## Discussion

The identification and quantification of the drivers of natural populations remains a long-standing question in ecology (Sutherland et al., 2013). Past efforts to tackle this challenge have focused on the temporal fluctuation of vital rates (*e.g.,* survival, reproduction; Morris et al., 2008; Sæther et al., 2016). Yet, the structure of a population (*i.e.,* the distribution of individuals among state classes such as age or stage) can also carry a legacy of the population’s recent history (Stott et al., 2011). This legacy generates transient (*i.e.,* short-term) dynamics that can account for half or more of the variation in population growth in variable environments (Ellis & Crone, 2013; McDonald et al., 2016). Whether that historical contribution to variation in growth rate can be predicted by life history traits, rather than having to be estimated *ad hoc* population by population, has remained unknown. Here, we simulate stochastic dynamics using empirical data across 137 plant and animal species, and partition the resulting variance in realised growth rate with an exact decomposition (Hernández et al., 2026) into contributions from vital-rate fluctuations and from transient dynamics to test the predictive power of life history traits. We find, contrary to theoretical expectations (Hernández et al., 2026), that populations with longer generation time have a smaller transient contribution to the variation in population growth rate. We also find that how survival rates vary along the life cycle of a population (survival unevenness) is a strong predictor of transient contribution in both plants and animals.

Generation time predicts ecological, evolutionary, and conservation properties alike. Examples include the timescale over which population declines are assessed (IUCN, 2024) to whether adaptation can keep pace with environmental change (Chevin et al., 2010). In the context of life history theory, one would expect the population dynamics of slow life histories (*i.e.*, those with long generation times) to be strongly affected by transient dynamics (Koons et al., 2005). This expectation reflects that the slow turnover typical of slow-living species causes structural departures from the stable distribution to decay only gradually (Hernández et al., 2026). Here, we find the reverse pattern in both plants and animals. We argue that, rather than contradicting theoretical expectations, our findings pinpoint key assumptions in the theory that do not perhaps hold in natural populations. The theoretical prediction from Hernández et al. (2026) assumes small shifts in vital rates due to annual variation in the environment, a condition rarely met in nature (Doak et al., 2005). By imposing large vital-rate fluctuations from real observations in our simulations, we find that faster life histories may be more transient-prone than slow ones. A possible explanation is that, in fast-lived populations, repeated shocks substantially redistribute individuals among rapidly turning-over state classes, preventing population structure from remaining close to its stable distribution. By contrast, the high adult survival characteristic of slow life histories may buffer the immediate structural effects of such shocks (Hilde et al., 2020; Morris et al., 2008). Whether transient dynamics scale with the pace of life history may therefore depend on the magnitude of environmental variation.

We identify survival unevenness as a strong predictor of the transient contribution to variation in population growth rate. Surprisingly, this effect was even stronger than that of generation time in plants. This finding is notable because comparative demography has long ranked life histories predominantly along the fast–slow continuum (Gaillard et al., 1989; Stearns, 1998), and a secondary axis defined by reproductive strategy or developmental pattern (Salguero-Gómez et al., 2016a; Stott et al., 2024), but neither of those two continua explicitly capture how survival is distributed across the life cycle of species. Survival unevenness has drawn less attention in comparative demography, although Healy et al. (2019) found that a similar metric explains substantial life history variation in animals in a deterministic environment. Our result provides a clear functional role to this otherwise underused life history trait, extending recent plant-based evidence that life history governs the demographic importance of population structure (Cant et al., 2025): survival unevenness predicts when a population’s structure shapes its growth in stochastic environments. As transient dynamics emerge whenever populations deviate from their stable stage distribution, the magnitude and direction of said dynamics are governed by the structure of the life cycle, which can transiently amplify rather than simply dampen these deviations (Neubert & Caswell, 1997; Stott et al., 2011). Uneven stage-specific survival along the life cycle of a species plausibly heightens these transient dynamics: when life cycle stages differ sharply in survival, they contribute unequally to the subsequent growth (Horvitz et al., 1997), so a given structural departure produces a larger swing in realised growth.

In plants, the populations with the most variable growth appear to also be those in which transient dynamics make the largest contribution to that variability. This relationship weakens to near zero after accounting for key life history traits and covariates like adult body size and length of the life stage. This result indicates that the predictors with the largest effect sizes (*i.e.*, survival unevenness and generation time) amplify temporal variation in population growth rate by generating stronger transient dynamics. Our findings carry a direct implication for management: conservation biology interventions typically target vital rates, for instance, the use of supplementary feeding to increase the survival and breeding of threatened birds (Fenn et al., 2020). However, where our results predict a large transient contribution, restoring or managing population structure may matter as much as improving survival or reproduction (Ezard et al., 2010). This is because an unfavourable structure can depress short-term growth and delay recovery even when vital rates are improved (Stott et al., 2011). Life history traits thus offer a rapid triage, indicating when a population’s departure from its stable structure, and not only its vital rates, should guide intervention, even where long demographic time series are unavailable (Capdevila et al., 2022; Jaggi et al., 2024). With the advent of large open-access demographic databases (*e.g.*, PADRINO, COMPADRE; Levin et al., 2022; Salguero-Gómez et al., 2015), life history can now inform conservation across species.

Our conclusions regarding how life history predicts the transient contribution are firmest in plants yet remain provisional in animals. The effect of iteroparity even reverses in sign between taxa: more iteroparous populations show a smaller transient contribution in plants but a larger one in animals. As such, we interpret this contrast with caution, given the marked difference in sample size between animals (n = 25 species) and plants (n = 112 species). Even the broadest animal demographic syntheses to date rest on only a few hundred species (Healy et al., 2019; Salguero-Gómez et al., 2016b), and demographic coverage remains sparse and taxonomically skewed (Conde et al., 2019); our own animal sample spans disparate taxa and offers little power to separate genuine differences from sampling noise. We therefore present the plant–animal contrasts as hypotheses for richer animal data to test, rather than as established biological facts.

Our stochastic simulations rest on the perturbations that populations have actually experienced in the wild, and disturbances that are novel in kind, or extreme in magnitude, may elicit different vital-rate responses. Here, we estimated the transient contribution under natural, unmanipulated conditions and tested how life history traits predict that contribution, so our approach samples only part of the perturbation space that ongoing change will impose (Koons et al., 2016). Beyond the perturbation magnitudes we sampled, vital-rate responses could be non-linear: rates that improve under moderate change can reverse under extreme conditions, so which rates respond, and how, need not remain fixed (Doak & Morris, 2010). Qualitatively novel disturbances raise a further difficulty, since they may act on rates that natural environmental fluctuation never samples (Johnstone et al., 2016). However, the matrix population models from experimental treatments now held in COMPADRE and COMADRE (n = 2,973) offer a direct way to test whether vital-rate responses to imposed disturbance differ from natural fluctuation.

Our framework would gain most from two advances: representing environments that change directionally, and deriving from first principles why particular life histories amplify transient dynamics. The first advance relaxes the assumption of stationarity (Wolkovich et al., 2014). Many pressures of global change intensify steadily over time, *i.e.*, ramp disturbance *sensu* Lake (2000), whereas our simulations resample the variation a population has already experienced and so hold the environmental regime fixed. Statistical methods can now reconstruct how directional environmental change reshapes vital rates from readily available data (González & Martorell, 2013), and building such environmentally explicit dynamics into the decomposition (Ehrlen et al., 2016) would extend prediction of the transient contribution to populations under sustained directional change, whose responses depend in turn on their life histories (Albaladejo-Robles et al., 2023). The second advance is mechanistic. Our trait associations are correlational, and grounding those associations between life history and transient contribution in mechanism would convert prediction into explanation, a progression that demographic buffering has already followed from pattern to process (Hilde et al., 2020).

By separating the demographic legacy carried by population structure from contemporaneous vital-rate changes, we show that the transient contribution to variation in population growth is indeed predictable from life history. Across our comparative dataset, transient contributions declined with generation time and increased with survival unevenness, with the clearest evidence in plants. These findings shift the question from whether transient dynamics matter to when they are most likely to shape population performance. Accounting for population structure alongside vital rates should therefore improve predictions of population responses to environmental change and bring the history-dependent side of population dynamics within the predictive reach of comparative life history theory.

## Supporting information

Supporting Imformation

## Author contributions

HL, CMH and RSG designed the study. HL and CMH performed the data collection, the analyses and wrote the associated code. HL wrote the first draft of the manuscript with input from RSG and WZ. All authors contributed substantially to revisions.

## Data accessibility

The data that support the findings of this study are openly available at www.compadre-db.org. All code files required to repeat the analyses are archived at Zenodo (https://doi.org/10.5281/zenodo.22100641).

## Notes

### Competing Interest Statement

The authors have declared no competing interest.

## Reference

Albaladejo-Robles, G., Böhm, M., & Newbold, T. (2023). Species life-history strategies affect population responses to temperature and land-cover changes. Global Change Biology, 29(1), 97–109.

Bienvenu, F., & Legendre, S. (2015). A new approach to the generation time in matrix population models. The American Naturalist, 185(6), 834–843.

Boer, G. J., Flato, G., & Ramsden, D. (2000). A transient climate change simulation with greenhouse gas and aerosol forcing: Projected climate to the twenty-first century. Climate Dynamics, 16(6), 427–450.

Botsford, L. W., Holland, M. D., Field, J. C., & Hastings, A. (2014). Cohort resonance: A significant component of fluctuations in recruitment, egg production, and catch of fished populations. ICES Journal of Marine Science, 71(8), 2158–2170.

Bürkner, P.-C. (2021). Bayesian item response modeling in R with brms and Stan. Journal of Statistical Software, 100, 1–54.

Cant, J. I., Hernandez, C. M., Koons, D., Hodgson, D. J., Qi, M., Hector, A., Stott, I., & Salguero-Gomez, R. (2025). Life-history variation mediates the importance of population structure to the short-term dynamics of plant populations worldwide.

Capdevila, P., Stott, I., Beger, M., & Salguero-Gómez, R. (2020). Towards a Comparative Framework of Demographic Resilience. Trends in Ecology & Evolution, 35(9), 776– 786. 10.1016/j.tree.2020.05.001

Capdevila, P., Stott, I., Cant, J., Beger, M., Rowlands, G., Grace, M., & Salguero-Gómez, R. (2022). Life history mediates the trade-offs among different components of demographic resilience. Ecology Letters, 25(6), Article 6. 10.1111/ele.14004

Caswell, H. (2001). Matrix population models. Sinauer Sunderland.

Chevin, L.-M., Lande, R., & Mace, G. M. (2010). Adaptation, plasticity, and extinction in a changing environment: Towards a predictive theory. PLoS Biology, 8(4), e1000357.

Cohen, J. E., Xu, M., & Schuster, W. S. (2012). Allometric scaling of population variance with mean body size is predicted from Taylor’s law and density-mass allometry. Proceedings of the National Academy of Sciences, 109(39), 15829–15834.

Conde, D. A., Staerk, J., Colchero, F., da Silva, R., Schöley, J., Baden, H. M., Jouvet, L., Fa, J. E., Syed, H., & Jongejans, E. (2019). Data gaps and opportunities for comparative and conservation biology. Proceedings of the National Academy of Sciences, 116(19), 9658–9664.

Crone, E. E., Menges, E. S., Ellis, M. M., Bell, T., Bierzychudek, P., Ehrlén, J., Kaye, T. N., Knight, T. M., Lesica, P., & Morris, W. F. (2011). How do plant ecologists use matrix population models? Ecology Letters, 14(1), 1–8.

Demetrius, L. (1978). Adaptive value, entropy and survivorship curves. Nature, 275(5677), 213–214.

Doak, D. F., & Morris, W. F. (2010). Demographic compensation and tipping points in climate-induced range shifts. Nature, 467(7318), 959–962.

Doak, D. F., Morris, W. F., Pfister, C., Kendall, B. E., & Bruna, E. M. (2005). Correctly estimating how environmental stochasticity influences fitness and population growth. The American Naturalist, 166(1), E14–E21.

Dunham, K. D., Tucker, A. M., Koons, D. N., Abebe, A., Dobson, F. S., & Grand, J. B. (2021). Demographic responses to climate change in a threatened Arctic species. Ecology and Evolution, 11(15), 10627–10643.

Ehrlen, J., Morris, W. F., von Euler, T., & Dahlgren, J. P. (2016). Advancing environmentally explicit structured population models of plants. Journal of Ecology, 104(2), 292–305.

Ellis, M. M., & Crone, E. E. (2013). The role of transient dynamics in stochastic population growth for nine perennial plants. Ecology, 94(8), 1681–1686.

Ellner, S. P., & Rees, M. (2006). Integral projection models for species with complex demography. The American Naturalist, 167(3), 410–428.

Ezard, T. H., Bullock, J. M., Dalgleish, H. J., Millon, A., Pelletier, F., Ozgul, A., & Koons, D. N. (2010). Matrix models for a changeable world: The importance of transient dynamics in population management. Journal of Applied Ecology, 47(3), 515–523.

Fenn, S. R., Bignal, E. M., Trask, A. E., McCracken, D. I., Monaghan, P., & Reid, J. M. (2020). Collateral benefits of targeted supplementary feeding on demography and growth rate of a threatened population. Journal of Applied Ecology, 57(11), 2212–2221.

Festa-Bianchet, M., Gaillard, J.-M., & Côté, S. D. (2003). Variable age structure and apparent density dependence in survival of adult ungulates. Journal of Animal Ecology, 72(4), 640–649.

Freckleton, R. P., Harvey, P. H., & Pagel, M. (2002). Phylogenetic analysis and comparative data: A test and review of evidence. The American Naturalist, 160(6), 712–726.

Gaillard, J.-M., Pontier, D., Allainé, D., Lebreton, J. D., Trouvilliez, J., & Clobert, J. (1989). An analysis of demographic tactics in birds and mammals. Oikos, 59–76.

Gaillard, J.-M., & Yoccoz, N. G. (2003). Temporal variation in survival of mammals: A case of environmental canalization? Ecology, 84(12), 3294–3306.

Gascoigne, S. J., Kajin, M., Tuljapurkar, S., Santos, G. S., Compagnoni, A., Steiner, U. K., Vinton, A. C., Jaggi, H., Sepil, I., & Salguero-Gómez, R. (2025). Structured demographic buffering: A framework to explore the environmental components and demographic mechanisms underlying demographic buffering. Ecology Letters, 28(2), e70066.

González, E. J., & Martorell, C. (2013). Reconstructing shifts in vital rates driven by long-term environmental change: A new demographic method based on readily available data. Ecology and Evolution, 3(7), 2273–2284.

Grant, A., & Benton, T. G. (1996). The impact of environmental variation on demographic convergence of Leslie matrix population models: An assessment using Lyapunov characteristic exponents. Theoretical Population Biology, 50(1), 18–30.

Healy, K., Ezard, T. H., Jones, O. R., Salguero-Gómez, R., & Buckley, Y. M. (2019). Animal life history is shaped by the pace of life and the distribution of age-specific mortality and reproduction. Nature Ecology & Evolution, 3(8), 1217–1224.

Hernández, C. M., Jaggi, H., Cant, J., Zuo, W., Tuljapurkar, S., & Salguero-Gomez, R. (2026). A robust method for quantifying the contribution of transient dynamics to variation in population growth rate.

Hilde, C. H., Gamelon, M., Sæther, B.-E., Gaillard, J.-M., Yoccoz, N. G., & Pélabon, C. (2020). The demographic buffering hypothesis: Evidence and challenges. Trends in Ecology & Evolution, 35(6), 523–538.

Horvitz, C., Schemske, D. W., & Caswell, H. (1997). The relative “importance” of life-history stages to population growth: Prospective and retrospective analyses. In Structured-population models in marine, terrestrial, and freshwater systems (pp. 247–271). Springer.

IUCN. (2024). Guidelines for Using the IUCN Red List Categories and Criteria. Version 16. IUCN Species Survival Commission.

Jaggi, H., Steinsaltz, D., & Tuljapurkar, S. (2024). Temporal variability can promote migration between habitats. Theoretical Population Biology, 158, 195–205.

Jaggi, H., Tuljapurkar, S., Zuo, W., Gascoigne, S. J., Kajin, M., & Salguero-Gómez, R. (2026). Transient dynamics and nonlinear fitness: A matrix approach to pulse and press perturbation. Ecology, 107(6), e70433.

Jenouvrier, S., Holland, M., Stroeve, J., Barbraud, C., Weimerskirch, H., Serreze, M., & Caswell, H. (2012). Effects of climate change on an emperor penguin population: Analysis of coupled demographic and climate models. Global Change Biology, 18(9), 2756–2770.

Jiang, S., Jaggi, H., Zuo, W., Oli, M. K., Coulson, T., Gaillard, J.-M., & Tuljapurkar, S. (2022). Reproductive dispersion and damping time scale with life-history speed. Ecology Letters, 25(9), 1999–2008.

Johnstone, J. F., Allen, C. D., Franklin, J. F., Frelich, L. E., Harvey, B. J., Higuera, P. E., Mack, M. C., Meentemeyer, R. K., Metz, M. R., & Perry, G. L. (2016). Changing disturbance regimes, ecological memory, and forest resilience. Frontiers in Ecology and the Environment, 14(7), 369–378.

Koons, D. N., Grand, J. B., Zinner, B., & Rockwell, R. F. (2005). Transient population dynamics: Relations to life history and initial population state. Ecological Modelling, 185(2–4), 283–297.

Koons, D. N., Iles, D. T., Schaub, M., & Caswell, H. (2016). A life-history perspective on the demographic drivers of structured population dynamics in changing environments. Ecology Letters, 19(9), 1023–1031.

Koons, D. N., Pavard, S., Baudisch, A., & Jessica E. Metcalf, C. (2009). Is life-history buffering or lability adaptive in stochastic environments? Oikos, 118(7), 972–980.

Lake, P. S. (2000). Disturbance, patchiness, and diversity in streams. Journal of the North American Benthological Society, 19(4), 573–592.

Lange, K. L., Little, R. J., & Taylor, J. M. (1989). Robust statistical modeling using the t distribution. Journal of the American Statistical Association, 84(408), 881–896.

Lemoine, N. P. (2019). Moving beyond noninformative priors: Why and how to choose weakly informative priors in Bayesian analyses. Oikos, 128(7), 912–928.

Levin, S. C., Evers, S., Potter, T., Guerrero, M. P., Childs, D. Z., Compagnoni, A., Knight, T. M., & Salguero-Gómez, R. (2022). Rpadrino: An R package to access and use PADRINO, an open access database of Integral Projection Models. Methods in Ecology and Evolution, 13(9), 1923–1929.

McDonald, J. L., Stott, I., Townley, S., & Hodgson, D. J. (2016). Transients drive the demographic dynamics of plant populations in variable environments. Journal of Ecology, 104(2), 306–314.

Morozov, A. Y., Almutairi, D., Petrovskii, S. V., & Hastings, A. (2024). Regime shifts, extinctions and long transients in models of population dynamics with density-dependent dispersal. Biological Conservation, 290, 110419.

Morris, W. F., Pfister, C. A., Tuljapurkar, S., Haridas, C. V., Boggs, C. L., Boyce, M. S., Bruna, E. M., Church, D. R., Coulson, T., & Doak, D. F. (2008). Longevity can buffer plant and animal populations against changing climatic variability. Ecology, 89(1), 19–25.

Neubert, M. G., & Caswell, H. (1997). Alternatives to resilience for measuring the responses of ecological systems to perturbations. Ecology, 78(3), 653–665.

Pagel, M. (1999). Inferring the historical patterns of biological evolution. Nature, 401(6756), 877–884.

R Core Team. (2020). RA language and environment for statistical computing, R Foundation for Statistical. Computing.

Räisänen, J. (2002). CO 2-induced changes in interannual temperature and precipitation variability in 19 CMIP2 experiments. Journal of Climate, 15(17), 2395–2411.

Sæther, B.-E., Grøtan, V., Engen, S., Coulson, T., Grant, P. R., Visser, M. E., Brommer, J. E., Rosemary Grant, B., Gustafsson, L., & Hatchwell, B. J. (2016). Demographic routes to variability and regulation in bird populations. Nature Communications, 7(1), 12001.

Salguero-Gómez, R., Jones, O. R., Archer, C. R., Bein, C., de Buhr, H., Farack, C., Gottschalk, F., Hartmann, A., Henning, A., & Hoppe, G. (2016). COMADRE: a global data base of animal demography. Journal of Animal Ecology, 85(2), 371–384.

Salguero-Gómez, R., Jones, O. R., Archer, C. R., Buckley, Y. M., Che-Castaldo, J., Caswell, H., Hodgson, D., Scheuerlein, A., Conde, D. A., & Brinks, E. (2015). The compadre P lant M atrix D atabase: An open online repository for plant demography. Journal of Ecology, 103(1), 202–218.

Salguero-Gómez, R., Jones, O. R., Jongejans, E., Blomberg, S. P., Hodgson, D. J., Mbeau-Ache, C., Zuidema, P. A., De Kroon, H., & Buckley, Y. M. (2016). Fast–slow continuum and reproductive strategies structure plant life-history variation worldwide. Proceedings of the National Academy of Sciences, 113(1), 230–235.

Schielzeth, H. (2010). Simple means to improve the interpretability of regression coefficients. Methods in Ecology and Evolution, 1(2), 103–113.

Sibly, R. M., Barker, D., Denham, M. C., Hone, J., & Pagel, M. (2005). On the regulation of populations of mammals, birds, fish, and insects. Science, 309(5734), 607–610.

Smallegange, I. M., & Berg, M. P. (2019). A functional trait approach to identifying life history patterns in stochastic environments. Ecology and Evolution, 9(16), 9350–9361.

Stearns, S. C. (1998). The evolution of life histories. Oxford university press.

Stott, I., Salguero-Gómez, R., Jones, O. R., Ezard, T. H., Gamelon, M., Lachish, S., Lebreton, J.-D., Simmonds, E. G., Gaillard, J.-M., & Hodgson, D. J. (2024). Life histories are not just fast or slow. Trends in Ecology & Evolution, 39(9), 830–840.

Stott, I., Townley, S., & Hodgson, D. J. (2011). A framework for studying transient dynamics of population projection matrix models. Ecology Letters, 14(9), 959–970.

Sutherland, W. J., Freckleton, R. P., Godfray, H. C. J., Beissinger, S. R., Benton, T., Cameron, D. D., Carmel, Y., Coomes, D. A., Coulson, T., & Emmerson, M. C. (2013). Identification of 100 fundamental ecological questions. Journal of Ecology, 101(1), 58–67.

Tenhumberg, B., Tyre, A. J., & Rebarber, R. (2009). Model complexity affects transient population dynamics following a dispersal event: A case study with pea aphids. Ecology, 90(7), 1878–1890.

Tuljapurkar, S. (1982). Population dynamics in variable environments. II. Correlated environments, sensitivity analysis and dynamics. Theoretical Population Biology, 21(1), 114–140.

Tuljapurkar, S. (1984). Demography in stochastic environments. I. Exact distributions of age structure. Journal of Mathematical Biology, 19(3), 335–350.

Tuljapurkar, S. (1990). Population dynamics in variable environments. Berlin, Heidelberg: Springer-Verlag.

Tuljapurkar, S., Gaillard, J.-M., & Coulson, T. (2009). From stochastic environments to life histories and back. Philosophical Transactions of the Royal Society B: Biological Sciences, 364(1523), 1499–1509.

Turner, M. G. (2010). Disturbance and landscape dynamics in a changing world. Ecology, 91(10), 2833–2849. 10.1890/10-0097.1

White, J. W., Botsford, L. W., Hastings, A., Baskett, M. L., Kaplan, D. M., & Barnett, L. A. (2013). Transient responses of fished populations to marine reserve establishment. Conservation Letters, 6(3), 180–191.

Williams, J. L., Ellis, M. M., Bricker, M. C., Brodie, J. F., & Parsons, E. W. (2011). Distance to stable stage distribution in plant populations and implications for near-term population projections. Journal of Ecology, 99(5), 1171–1178.

Wolkovich, E. M., Cook, B. I., McLauchlan, K. K., & Davies, T. J. (2014). Temporal ecology in the Anthropocene. Ecology Letters, 17(11), 1365–1379.

Zuur, A. F., Ieno, E. N., & Elphick, C. S. (2010). A protocol for data exploration to avoid common statistical problems. Methods in Ecology and Evolution, 1(1), 3–14.

