## Supporting Imformation for "Life history traits predict the contribution of transient dynamics to variation in population growth"

Contents


| | |
|---|---|
| Supplementary Table 1 | 2 |
| Supplementary Table 2 | 6 |
| Supplementary Table 3 | 7 |
| Supplementary Table 4 | 8 |
| Supplementary Figure 1 | 9 |
| Supplementary Figure 2 | 10 |
| Supplementary Figure 3 | 11 |
| Supplementary Figure 4 | 12 |
| Supplementary Figure 5 | 13 |
| Supplementary Figure 6 | 14 |
| Supplementary Figure 7 | 15 |
| Supplementary Figure 8 | 16 |
| Supplementary Methods | 17 |
| Reference | 21 |

**Supplementary Table 1. Body-size data and their sources for the 137 species analysed (112 plants and 25 animals).** Body size denotes maximum height (m) for plants and adult body mass (g) for animals, as indicated by the Unit column. Plant heights were drawn primarily from the TRY plant trait database and animal body masses from the MOSAIC database; where these sources lacked a value, body sizes were compiled in this study from floras, species monographs and herbarium accounts (plants) or peer-reviewed publications (animals), the latter including colony dry mass for the two coral species and a length–weight conversion for one fish species. The "Body size source" column records the origin of each value.

| Species | Body size | Unit | Body size source |
|---|---|---|---|
| *Alces alces* | 351000 | g | MOSAIC |
| *Anthropoides paradiseus* | 6200 | g | del Hoyo, J. et al. (1996) |
| *Brachyteles hypoxanthus* | 10840 | g | MOSAIC |
| *Callospermophilus lateralis* | 340 | g | Ferron, J. (1985) |
| *Cebus capucinus* | 2655 | g | MOSAIC |
| *Centrocercus minimus* | 1650 | g | MOSAIC |
| *Cercopithecus mitis* | 6117 | g | MOSAIC |
| *Chen caerulescens* | 2631 | g | MOSAIC |
| *Clinocottus analis* | 10 | g | FishBase |
| *Falco peregrinus* | 770 | g | Dunning, J. B. (2008) |
| *Gorilla beringei beringei* | 110000 | g | Williamson, E. A., & Butynski, T. M. (2013) |
| *Macaca mulatta* | 6614 | g | MOSAIC |
| *Orcinus orca* | 4300000 | g | MOSAIC |
| *Ovis aries* | 110000 | g | MOSAIC |
| *Pan troglodytes schweinfurthii* | 37000 | g | Pusey, A. E. et al. (2005) |
| *Papio cynocephalus* | 17851 | g | MOSAIC |
| *Paramuricea clavata* | 220 | g | Rossi, S., et al. (2017) |
| *Propithecus verreauxi* | 3588 | g | MOSAIC |
| *Sceloporus grammicus* | 8 | g | Ramírez-Bautista, A., et al. (2012) |
| *Strix occidentalis* | 580 | g | Dunning, J. B. (2008) |
| *Suricata suricatta* | 731 | g | Jones, K. E., et al. (2009) |
| *Umbonium costatum* | 9 | g | Noda, T., & Nakao, S. (1998) |
| *Ursus maritimus* | 371704 | g | MOSAIC |
| *Xenosaurus grandis* | 24 | g | MOSAIC |
| *Xenosaurus platyceps* | 20 | g | MOSAIC |
| *Abies concolor* | 60 | m | TRY |
| *Abies magnifica* | 60 | m | TRY |
| *Actaea elata* | 2 | m | Burke Museum |
| *Actaea spicata* | 0.8 | m | TRY |
| *Adenocarpus gibbsianus* | 3 | m | Junta de Andalucía |
| *Agrimonia eupatoria* | 1.5 | m | TRY |
| *Alliaria petiolata* | 1.2 | m | TRY |

| | | | |
|---|---|---|---|
| *Alnus incana subsp. rugosa* | 4.8768 | m | TRY |
| *Anarrhinum fruticosum* | 1 | m | En el ecotono |
| *Ardisia elliptica* | 6 | m | TRY |
| *Arenaria grandiflora subsp. bolosii* | 0.15 | m | TRY |
| *Armeria merinoi* | 0.18 | m | Flora Vascular |
| *Asplenium adulterinum* | 0.2 | m | TRY |
| *Asplenium cuneifolium* | 0.45 | m | TRY |
| *Astragalus alopecurus* | 1 | m | TRY |
| *Astragalus peckii* | 0.3 | m | Oregon Department of Agriculture |
| *Astragalus scaphoides* | 0.4 | m | Atlas of North American Astragalus |
| *Astragalus tremolsianus* | 0.1 | m | Hoseito, Flores Silvestres de España |
| *Astragalus tyghensis* | 0.55 | m | Oregon Department of Agriculture |
| *Boechera fecunda* | 0.3 | m | Flora of North America |
| *Brassavola cucullata* | 0.2 | m | Smithsonian Institution Living Collections |
| *Brassica insularis* | 1 | m | Flora della Sicilia |
| *Calathea ovandensis* | 0.9 | m | Estimated from cultivated records of congeneric species |
| *Calocedrus decurrens* | 72 | m | TRY |
| *Calochortus lyallii* | 0.5 | m | Flora of North America |
| *Castanea dentata* | 35.052 | m | TRY |
| *Chamaecrista lineata var. keyensis* | 0.76 | m | U.S. Fish & Wildlife Service |
| *Cheirolophus metlesicsii* | 0.8 | m | TRY |
| *Cirsium pitcheri* | 1 | m | Center for Plant Conservation |
| *Cleistesiopsis bifaria* | 0.38 | m | USDA Forest Service |
| *Clidemia hirta* | 0.623 | m | TRY |
| *Corallorhiza trifida* | 0.3 | m | TRY |
| *Cypripedium calceolus* | 0.5 | m | TRY |
| *Cypripedium fasciculatum* | 0.2 | m | Columbia River Gorge flora |
| *Dicerandra frutescens* | 0.5 | m | Center for Plant Conservation |
| *Dioon merolae* | 6 | m | Whitelock (2002) |
| *Dracocephalum austriacum* | 0.4 | m | TRY |
| *Echinospartum ibericum subsp. algibicum* | 0.4 | m | TRY |
| *Eriogonum longifolium var. gnaphalifolium* | 0.4 | m | TRY |
| *Erodium paularense* | 0.2 | m | Flora de Aragón |
| *Eryngium alpinum* | 1 | m | TRY |
| *Eryngium cuneifolium* | 0.9 | m | Flora of North America |
| *Escobaria robbinsorum* | 0.06 | m | Flora of North America |
| *Euphorbia fontqueriana* | 0.15 | m | Ministerio para la Transición Ecológica y el Reto Demográfico |
| *Frasera speciosa* | 0.8 | m | TRY |
| *Gentiana pneumonanthe* | 8 | m | TRY |
| *Helianthemum juliae* | 0.57 | m | TRY |
| *Helianthemum polygonoides* | 0.57 | m | TRY |

| | | | |
|---|---|---|---|
| *Helianthemum teneriffae* | 0.57 | m | TRY |
| *Horkelia congesta* | 0.5 | m | Flora of North America |
| *Hypericum cumulicola* | 0.7 | m | Center for Plant Conservation |
| *Jurinea fontqueri* | 0.6 | m | TRY |
| *Kosteletzkya pentacarpos* | 1.8 | m | Flora of the Southeastern United States |
| *Laserpitium longiradium* | 1.8 | m | Atlas y Libro Rojo de la Flora Vascular Amenazada de España |
| *Lathyrus vernus* | 0.5 | m | TRY |
| *Lechea cernua* | 0.3 | m | Florida Natural Areas Inventory |
| *Lechea deckertii* | 0.3 | m | Flora of North America |
| *Lepidium davisii* | 0.14 | m | Flora of North America |
| *Liatris scariosa* | 1.5 | m | NC State Extension |
| *Limonium erectum* | 0.2 | m | TRY |
| *Limonium geronense* | 0.2 | m | TRY |
| *Limonium malacitanum* | 0.2 | m | TRY |
| *Linnaea borealis* | 10 | m | TRY |
| *Linum flavum* | 0.5 | m | TRY |
| *Linum tenuifolium* | 0.45 | m | TRY |
| *Lomatium bradshawii* | 0.65 | m | United States Department of Agriculture |
| *Lomatium cookii* | 0.5 | m | U.S. Fish & Wildlife Service |
| *Lotus arinagensis* | 0.1 | m | TRY |
| *Lupinus lepidus* | 1.8288 | m | TRY |
| *Lupinus tidestromii* | 0.3 | m | USFWS Sacramento Fish & Wildlife Office |
| *Mammillaria hernandezii* | 0.04 | m | Glass & Foster (1983) |
| *Mammillaria huitzilopochtli* | 0.15 | m | Hunt (1979) |
| *Mimulus cardinalis* | 0.762 | m | TRY |
| *Mimulus lewisii* | 1.07 | m | USDA Forest Service |
| *Molinia caerulea* | 2.5 | m | TRY |
| *Oenothera deltoides* | 0.46 | m | U.S. Fish & Wildlife Service |
| *Opuntia rastrera* | 0.6 | m | Promesse de Fleurs |
| *Orchis purpurea* | 1 | m | TRY |
| *Oxytropis jabalambrensis* | 0.3 | m | TRY |
| *Parolinia glabriuscula* | 0.8 | m | TRY |
| *Pediocactus bradyi* | 0.06 | m | Flora of North America |
| *Petrocoptis pyrenaica subsp. pseudoviscosa* | 0.3 | m | TRY |
| *Phyllanthus emblica* | 14 | m | TRY |
| *Pinus lambertiana* | 82 | m | TRY |
| *Pinus strobus* | 80 | m | TRY |
| Plantago coronopus | 0.3 | m | TRY |
| Plantago media | 0.8 | m | TRY |
| Polemonium van-bruntiae | 1.4 | m | Committee on the Status of Endangered Wildlife in Canada |
| Polygonella basiramia | 0.8 | m | Flora of North America |

| | | | |
|---|---|---|---|
| Primula elatior | 0.3 | m | TRY |
| Primula farinosa | 0.2 | m | TRY |
| Prosopis glandulosa | 12 | m | TRY |
| Purshia subintegra | 0.75 | m | Flora of North America |
| Pyrrocoma radiata | 1 | m | Oregon Department of Agriculture |
| Ramonda myconi | 0.1 | m | Royal Horticultural Society |
| Ranunculus peltatus | 2 | m | TRY |
| Rosmarinus tomentosus | 0.5 | m | Diputación de Málaga |
| Rumex rupestris | 0.7 | m | TRY |
| Santolina melidensis | 0.6 | m | TRY |
| Sapium sebiferum | 20 | m | TRY |
| Saponaria bellidifolia | 0.5 | m | Flore de la France |
| Silene acaulis | 0.1 | m | TRY |
| Silene spaldingii | 0.4 | m | U.S. Fish & Wildlife Service |
| Succisa pratensis | 1 | m | TRY |
| Taxus floridana | 9 | m | TRY |
| Trillium grandiflorum | 0.5 | m | TRY |
| Trillium ovatum | 0.5 | m | Flora of North America |
| Trollius europaeus | 0.845 | m | TRY |
| Trollius laxus | 0.29 | m | TRY |
| Vella pseudocytisus subsp. paui | 2 | m | Domínguez-Lozano, F. et al. (2005) |
| Verticordia staminosa | 1 | m | Gardner, C. A., & George, A. S. (1963) |
| Zea diploperennis | 2.5 | m | Iltis et al. (1979) |

**Supplementary Table 2. Variance inflation factors quantifying collinearity among the predictors of the full multivariate phylogenetic models.** Variance inflation factors (VIFs) are shown separately for plants and animals, for the three life-history traits — generation time (*T*), degree of iteroparity (*S*), and survival unevenness (*Vσ*) — and the two covariates — body size (plant height or animal body mass, log-transformed) and matrix dimension. Values below 3 indicate low collinearity among predictors (Zuur et al., 2010); all values fall below this threshold except generation time in animals (VIF = 3.04), which is only marginally above it, indicating that collinearity is low throughout.

| Taxa | Predictor | Variance inflation factor |
|---|---|---|
| Plant | Generation time *T* | 1.09 |
| | Degree of iteroparity *S* | 1.14 |
| | Survival unevenness *Vσ* | 1.15 |
| | Plant height (log) | 1.07 |
| | Matrix dimension | 1.14 |
| Animal | Generation time *T* | 3.04 |
| | Degree of iteroparity *S* | 1.43 |
| | Survival unevenness *Vσ* | 1.95 |
| | Body mass (log) | 1.69 |
| | Matrix dimension | 2.88 |

**Supplementary Table 3. Posterior summaries of fixed effects from the full multivariate phylogenetic models.** Values are posterior means (Estimate), posterior standard deviations (S.D.), and 95% credible intervals for the effects of three life-history traits — generation time ($T$), degree of iteroparity ($S$), and survival unevenness ($V\sigma$) — and two covariates — body size (plant height or animal body mass, log-transformed) and matrix dimension — on two jointly modelled responses: temporal variation in population growth rate (log Var $\lambda_t$) and the proportional contribution of transient dynamics to that variation. Models were fitted separately for plants and animals, and included species identity (with phylogenetic covariance) and the number of contributing matrices as random effects, with Student-t response distributions. All predictors were standardised to zero mean and unit variance, so estimates are expressed in units of standard deviation and are comparable across traits. Coefficients whose 95% credible interval excludes zero are shown in bold.

| Taxa | Response | Predictor | Estimate | SD | 95% CI |
|---|---|---|---|---|---|
| Plant | Temporal variation in growth rate (log) | Intercept | −1.31 | 1.45 | [−4.19, 1.52] |
| | | Generation time *T* | **−1.76** | 0.27 | [−2.25, −1.23] |
| | | Degree of iteroparity *S* | **−0.48** | 0.11 | [−0.70, −0.26] |
| | | Survival unevenness *Vσ* | **0.39** | 0.15 | [0.10, 0.69] |
| | | Plant height (log) | **0.52** | 0.25 | [0.04, 1.01] |
| | | Matrix dimension | **0.47** | 0.20 | [0.06, 0.87] |
| | Contribution of transient dynamics (proportion) | Intercept | 0.40 | 0.23 | [−0.06, 0.86] |
| | | Generation time *T* | −0.06 | 0.03 | [−0.11, 0.001] |
| | | Degree of iteroparity *S* | **−0.03** | 0.01 | [−0.06, −0.003] |
| | | Survival unevenness *Vσ* | **0.12** | 0.02 | [0.08, 0.15] |
| | | Plant height (log) | −0.02 | 0.03 | [−0.08, 0.04] |
| | | Matrix dimension | 0.03 | 0.02 | [−0.02, 0.07] |
| Animal | Temporal variation in growth rate (log) | Intercept | **−3.32** | 1.12 | [−5.28, −0.92] |
| | | Generation time *T* | **−1.45** | 0.57 | [−2.53, −0.30] |
| | | Degree of iteroparity *S* | −0.42 | 0.43 | [−1.27, 0.40] |
| | | Survival unevenness *Vσ* | −0.14 | 0.41 | [−0.92, 0.71] |
| | | Body mass (log) | −0.07 | 0.55 | [−1.15, 1.01] |
| | | Matrix dimension | 0.11 | 0.53 | [−0.99, 1.14] |
| | Contribution of transient dynamics (proportion) | Intercept | **0.13** | 0.04 | [0.07, 0.21] |
| | | Generation time *T* | **−0.08** | 0.04 | [−0.16, −0.002] |
| | | Degree of iteroparity *S* | **0.06** | 0.03 | [0.01, 0.11] |
| | | Survival unevenness *Vσ* | 0.05 | 0.03 | [−0.004, 0.11] |
| | | Body mass (log) | 0.01 | 0.04 | [−0.05, 0.09] |
| | | Matrix dimension | 0.01 | 0.04 | [−0.07, 0.08] |

**Supplementary Table 4. Posterior summaries of fixed effects from the supplementary models that additionally include vital-rate variability (the coefficient of variation of each population's most influential vital rate) as a covariate.** Model structure, responses, random effects, and predictor standardisation follow Supplementary Table 2. Values are posterior means (Estimate), posterior standard deviations (S.D.), and 95% credible intervals; coefficients whose 95% credible interval excludes zero are shown in bold.

| Taxa | Response | Predictor | Estimate | SD | 95% CI |
|---|---|---|---|---|---|
| Plant | Temporal variation in growth rate (log) | Intercept | −1.33 | 1.40 | [−4.06, 1.44] |
| | | Generation time *T* | **−1.70** | 0.27 | [−2.21, −1.15] |
| | | Degree of iteroparity *S* | **−0.47** | 0.11 | [−0.69, −0.25] |
| | | Survival unevenness *V*σ | **0.42** | 0.15 | [0.12, 0.72] |
| | | Vital-rate variability | 0.19 | 0.12 | [−0.04, 0.43] |
| | | Plant height (log) | **0.49** | 0.24 | [0.04, 0.96] |
| | | Matrix dimension | **0.44** | 0.20 | [0.04, 0.84] |
| | Contribution of transient dynamics (proportion) | Intercept | 0.39 | 0.23 | [−0.07, 0.85] |
| | | Generation time *T* | −0.05 | 0.03 | [−0.11, 0.004] |
| | | Degree of iteroparity *S* | **−0.03** | 0.01 | [−0.06, −0.002] |
| | | Survival unevenness *V*σ | **0.11** | 0.02 | [0.08, 0.15] |
| | | Vital-rate variability | 0.00 | 0.01 | [−0.03, 0.03] |
| | | Plant height (log) | –0.02 | 0.03 | [−0.08, 0.04] |
| | | Matrix dimension | 0.03 | 0.02 | [−0.02, 0.07] |
| Animal | Temporal variation in growth rate (log) | Intercept | **−3.23** | 1.15 | [−5.22, −0.80] |
| | | Generation time *T* | **−1.25** | 0.59 | [−2.40, −0.08] |
| | | Degree of iteroparity *S* | −0.44 | 0.41 | [−1.25, 0.36] |
| | | Survival unevenness *V*σ | –0.19 | 0.39 | [−0.92, 0.62] |
| | | Vital-rate variability | 0.59 | 0.42 | [−0.26, 1.43] |
| | | Body mass (log) | 0.08 | 0.56 | [−1.02, 1.18] |
| | | Matrix dimension | 0.14 | 0.51 | [−0.91, 1.11] |
| | Contribution of transient dynamics (proportion) | Intercept | **0.13** | 0.04 | [0.06, 0.22] |
| | | Generation time *T* | −0.08 | 0.04 | [−0.17, 0.003] |
| | | Degree of iteroparity *S* | **0.06** | 0.03 | [0.01, 0.11] |
| | | Survival unevenness *V*σ | 0.05 | 0.03 | [−0.01, 0.11] |
| | | Vital-rate variability | −0.01 | 0.03 | [−0.07, 0.05] |
| | | Body mass (log) | 0.01 | 0.04 | [−0.06, 0.09] |
| | | Matrix dimension | 0.00 | 0.04 | [−0.07, 0.08] |

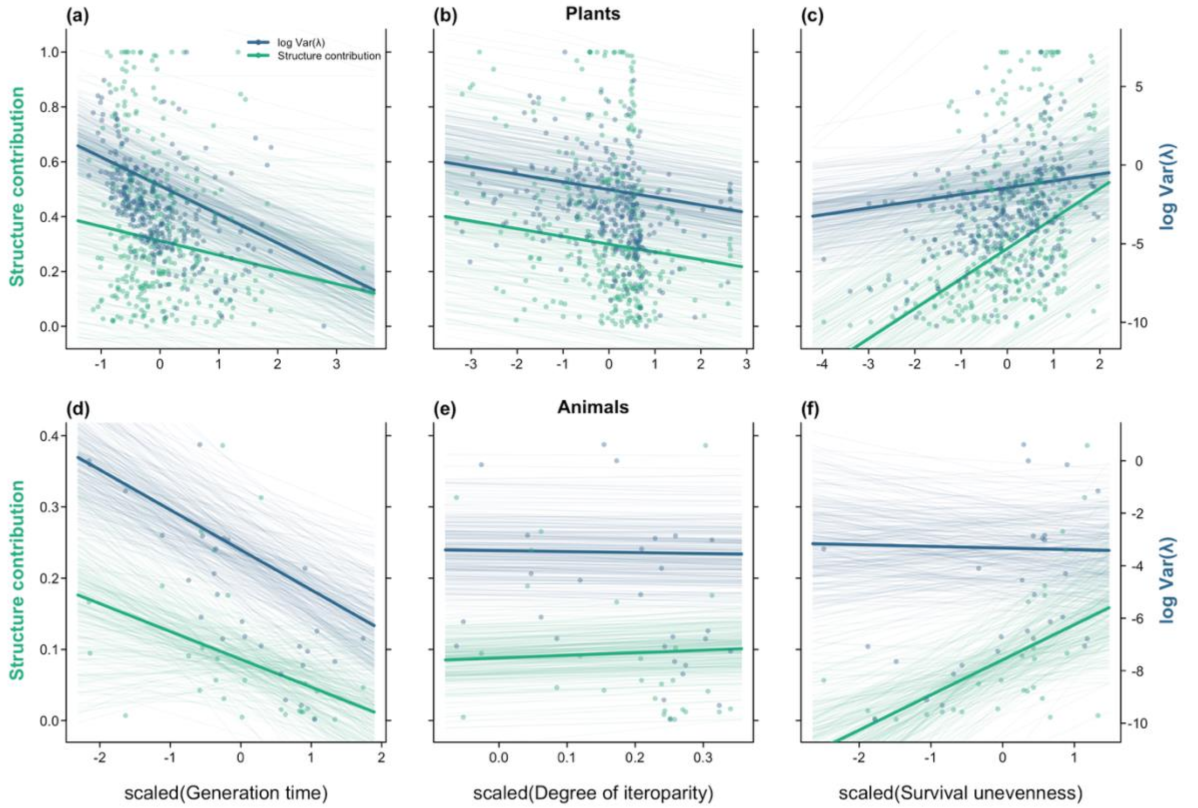


**Supplementary Figure 1. Life history predicts the population-structure component of the transient contribution as it does the full transient contribution.** Each panel relates one life history trait (x-axis) to the population-structure component of the transient contribution (Structure contribution; left y-axis, green) and log-transformed temporal variation in population growth rate (log Var(λ); right y-axis, blue), for plants (a–c) and animals (d–f). Panels show generation time (a, d), degree of iteroparity (b, e) and survival unevenness (c, f). All three life history traits were standardised to zero mean and unit variance, and each point represents one population. Thin lines show 250 draws from the posterior of the full phylogenetic multilevel model, and thick lines the posterior mean. For visual clarity, populations with extreme standardised trait values ($|z| > 5$) are omitted from the plotted points and axis ranges, but were retained in all analyses.

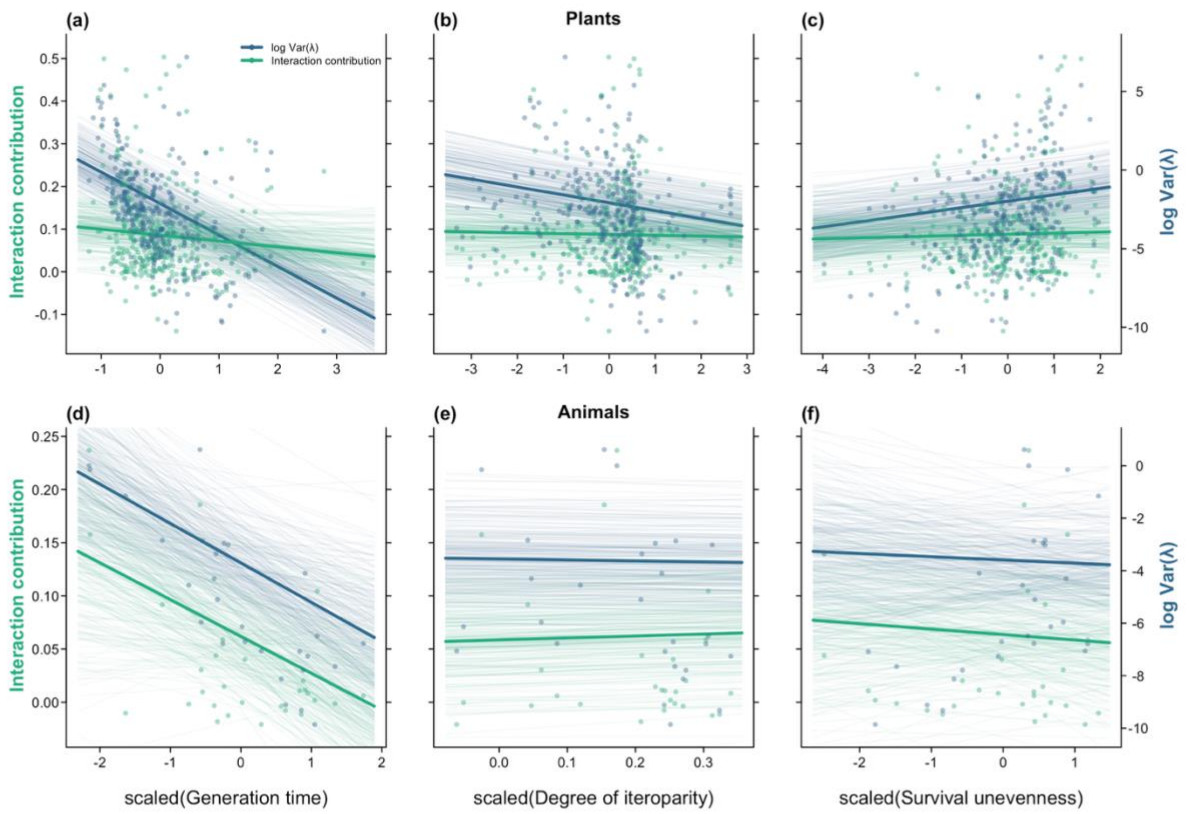


**Supplementary Figure 2. The interaction component of the transient contribution shows association only with generation time.** Each panel relates one life history trait (x-axis) to the interaction component of the transient contribution (Interaction contribution; left y-axis, green) and log-transformed temporal variation in population growth rate (log Var(λ); right y-axis, blue), for plants (a–c) and animals (d–f). Panels show generation time (a, d), degree of iteroparity (b, e) and survival unevenness (c, f). All three life history traits were standardised to zero mean and unit variance, and each point represents one population. Thin lines show 250 draws from the posterior of the full phylogenetic multilevel model, and thick lines the posterior mean. For visual clarity, populations with extreme standardised trait values (|z| > 5) are omitted from the plotted points and axis ranges, but were retained in all analyses.

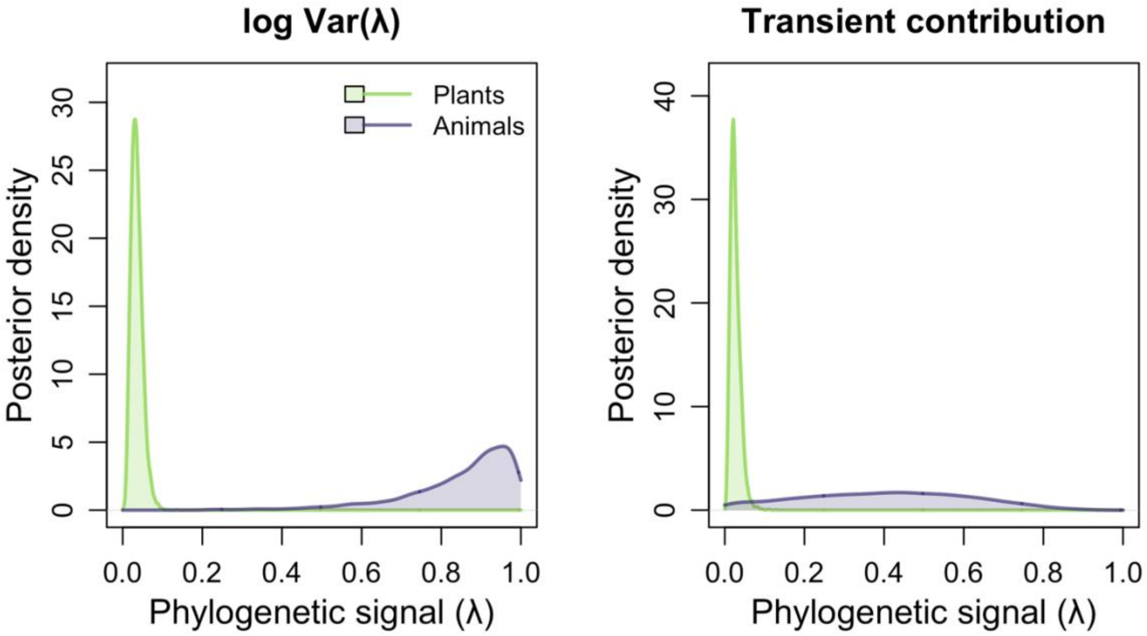


**Supplementary Figure 3. Phylogenetic signal in both response variables is negligible in plants but highly uncertain in animals.** Plants show negligible phylogenetic signal for both response variables (log-transformed temporal variance in population growth rate and the proportional contribution of transient dynamics to that variation), whereas animals show highly uncertain signal. Posterior distributions of Pagel's λ analogue — the proportion of total variance attributable to phylogenetic covariance — estimated from intercept-only phylogenetic multilevel models. Left and right panels show distributions for log-transformed temporal variation in population growth rate (log Var(λ)) and for the proportional contribution of transient dynamics to that variation (transient contribution), respectively. Green and purple shading indicate plants and animals, respectively. For plants, posterior mass is concentrated near zero for both response variables. For animals, distributions are broad and span the full parameter range, reflecting high uncertainty in phylogenetic signal estimates.

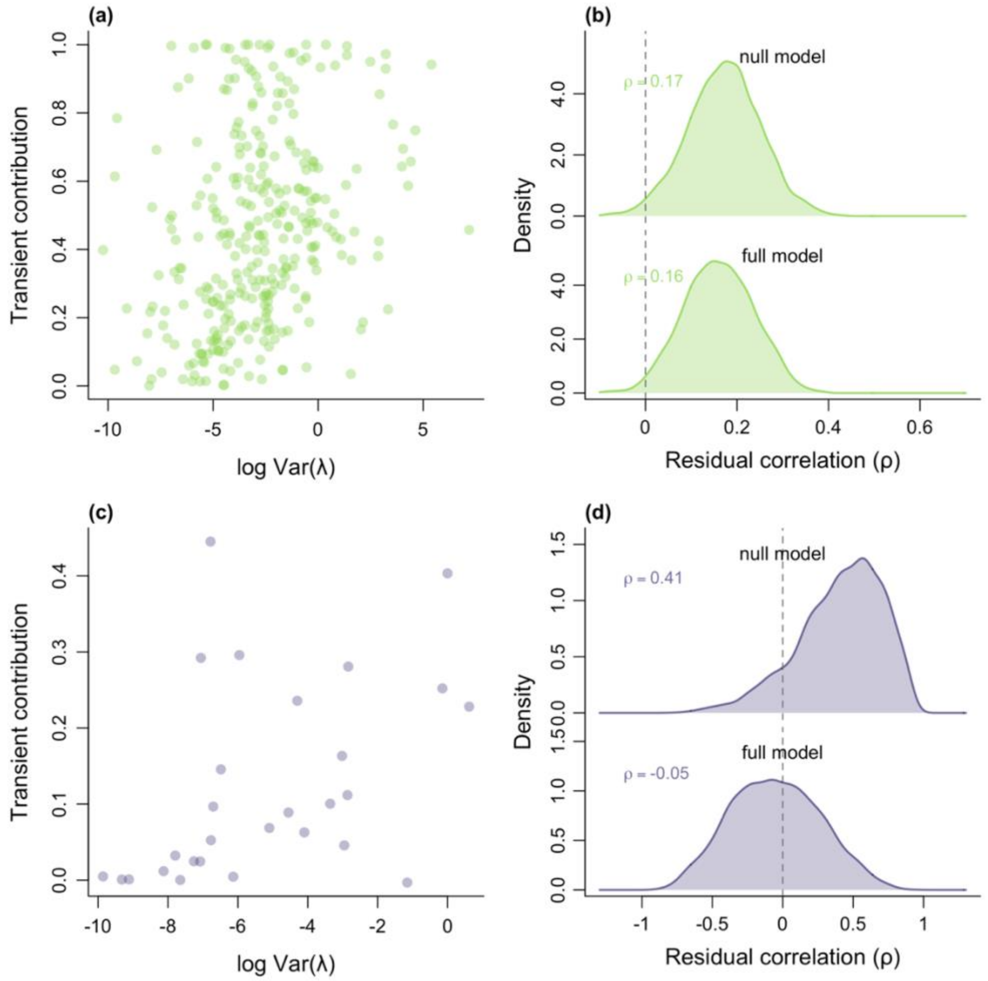


**Supplementary Figure 4. Without phylogenetic correction, both response variables show positive residual co-variation under the null model in both plants and animals.** Residual co-variation between the two response variables (log-transformed temporal variance in population growth rate and the proportional contribution of transient dynamics to that variation) in plants and animals, with and without predictor traits. (a, c) Scatter plots of log-transformed temporal variance in population growth rate (log Var(λ); x-axis) against the proportional contribution of transient dynamics to that variation (transient contribution; y-axis) for plants (a; green) and animals (c; purple). Each point represents one population. (b, d) Posterior density of the residual correlation (ρ) between the two response variables from the null model (intercept only; top) and the full model (with life-history predictors; bottom), for plants (b) and animals (d). The dashed vertical line indicates ρ = 0. In both taxa, the two response variables show positive residual co-variation under the null model. Adding the life-history predictors removes this co-variation in animals but not in plants (full ρ = 0.16, [0.01, 0.31]), where it is resolved only once phylogeny is also accounted for (Fig. 3).

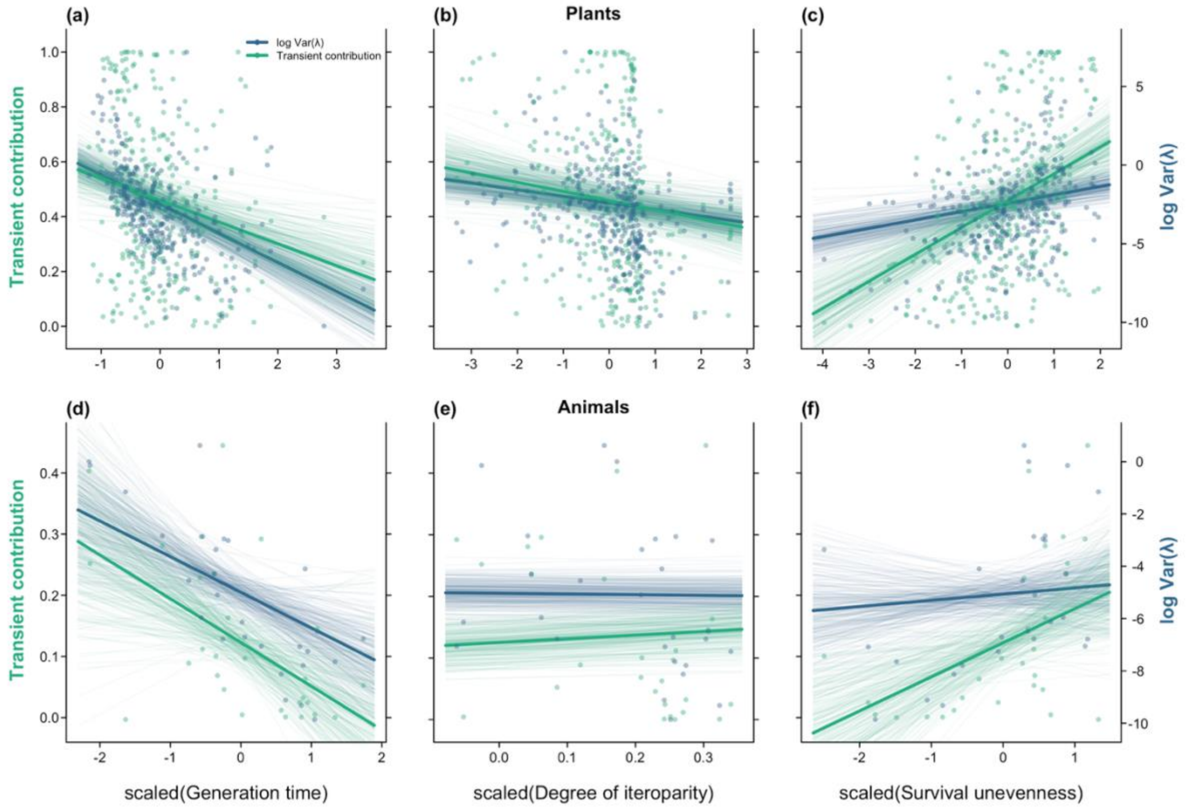


**Supplementary Figure 5. The associations between life history traits and both response variables are qualitatively consistent regardless of whether phylogenetic structure is accounted for.** Life history trait effects on both response variables ((log-transformed temporal variance in population growth rate and the proportional contribution of transient dynamics to that variation) are consistent between phylogenetically corrected and uncorrected analyses. Scatter plots show scaled life history traits on the x-axes against the proportional contribution of transient dynamics to the temporal variation in population growth rate (transient contribution; left y-axis, green), and the log-transformed temporal variation in population growth rate (log Var(λ); right y-axis, blue), for plants (panels a–c) and animals (panels d–f). Estimates are from a non-phylogenetic multivariate Bayesian model. Each point represents one population. Thin lines show 250 posterior draws; thick lines indicate posterior means. For visual clarity, three populations with extreme standardised trait values (|z| > 5; one each in panels a, b and e) are omitted from the plotted points and axis ranges; these populations were retained in all analyses, and the fitted lines shown reflect the full data. Left y-axes show transient contribution, and right y-axes show log Var(λ), with tick labels shown only on the rightmost panels (c, f).

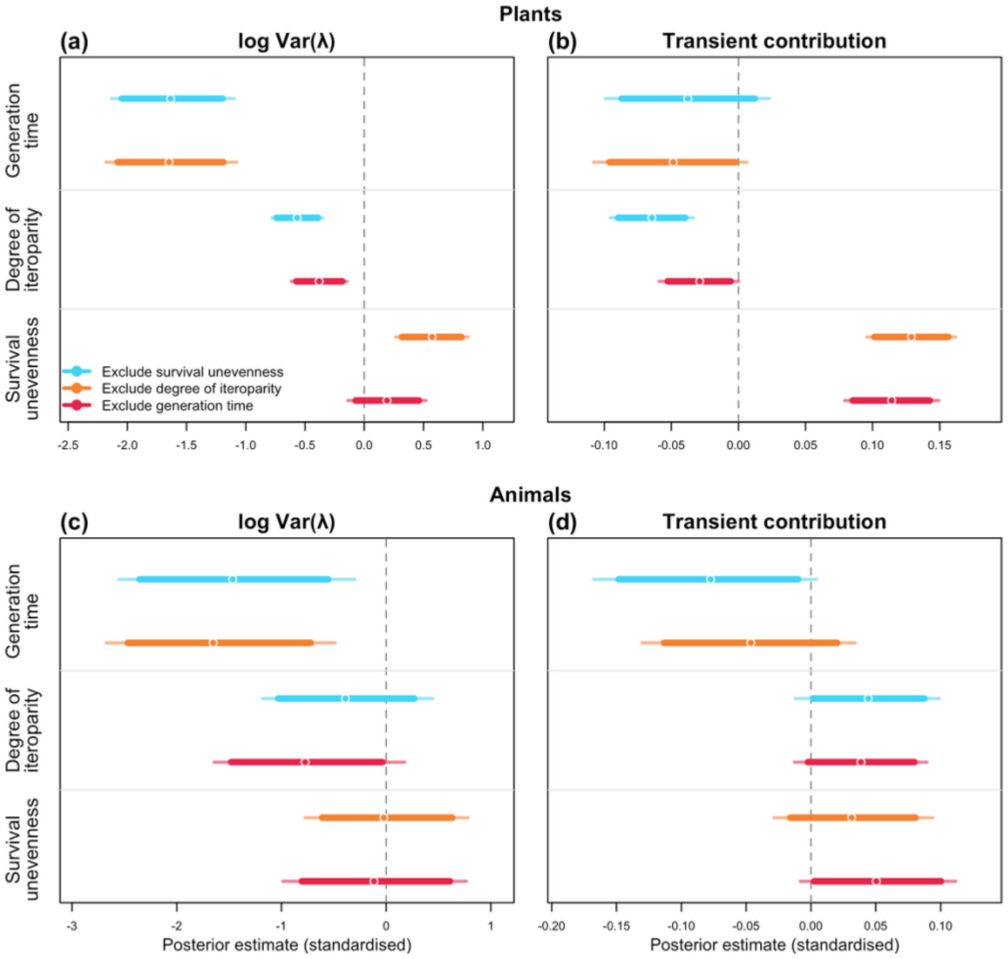


**Supplementary Figure 6. The associations between life history traits and response variables are robust to leaving out each focal trait in turn, in both plants and animals.** Leave-one-out sensitivity analysis in which each full phylogenetic multilevel model was refitted three times, each time excluding one of the three focal life-history traits (generation time, degree of iteroparity, and survival unevenness) while retaining all covariates (body size and matrix dimension). Panels show the posterior estimates (standardised; x-axis) of each life-history trait (y-axis) on the two jointly modelled responses: the log-transformed temporal variation in population growth rate (log Var(λ); panels a, c) and the proportional contribution of transient dynamics to that variation (transient contribution; panels b, d), for plants (a, b) and animals (c, d). Within each panel, the estimate for a given trait is shown once for every model in which that trait was retained (i.e., the two models that excluded one of the other two traits), coloured by which trait was excluded (blue, survival unevenness; orange, degree of iteroparity; red, generation time; legend in panel a). Points are posterior medians, thick bars are 89% credible intervals, and thin bars are 95% credible intervals; the dashed vertical line marks zero. Across the leave-one-out models, estimates are essentially unchanged for every trait and response, indicating the associations are not artefacts of collinearity. The one exception is survival unevenness in plants: its effect on growth-rate variation stays positive but weakens and overlaps zero when generation time is excluded, while its effect on the transient contribution remains stable.

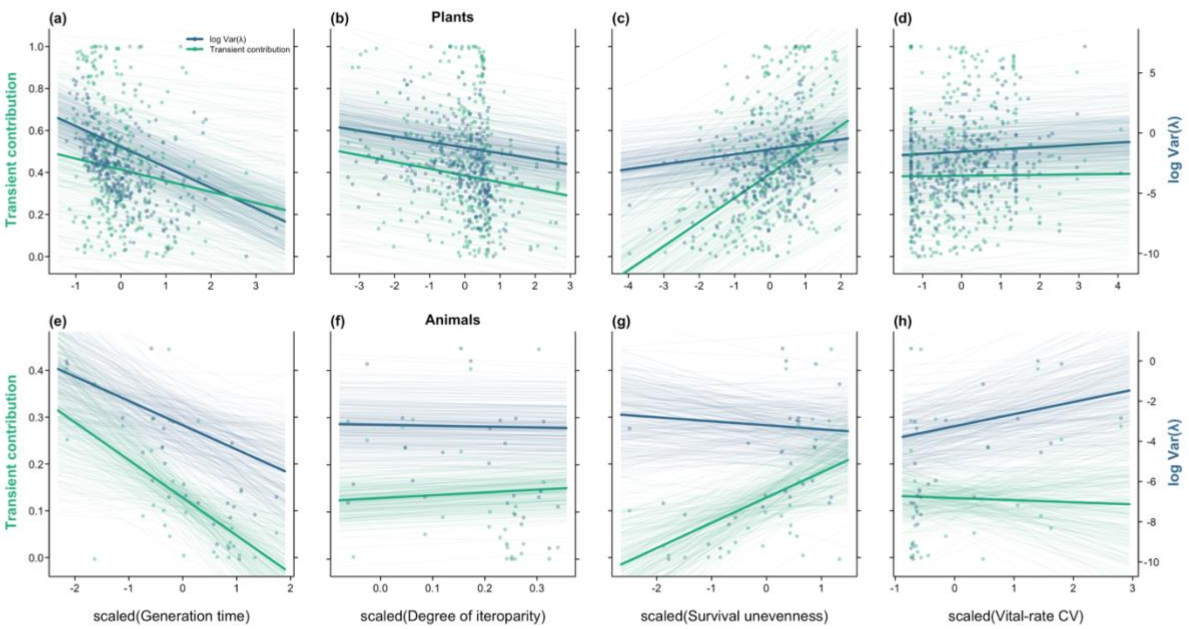


**Supplementary Figure 7. The associations of generation time, iteroparity, and survival unevenness with both response variables are qualitatively unchanged when vital-rate CV is included as an additional predictor.** Life history trait effects on both response variables ((log-transformed temporal variance in population growth rate and the proportional contribution of transient dynamics to population growth rate) are consistent with and without vital-rate CV (the temporal coefficient of variation of the most growth-sensitive matrix element) and vital-rate CV itself shows at most a weak, uncertain positive association with temporal variation in growth rate in plants (95% credible interval overlapping zero) and none with the transient contribution. Scatter plots of scaled life history traits (x-axes) against the proportional contribution of transient dynamics to population growth rate (transient contribution; left y-axis; green) and log-transformed temporal variation in population growth rate (log Var(λ); right y-axis; blue) for plants (panels a–d) and animals (panels e–h). Each point represents one population. Thin lines show 250 posterior draw lines sampled from the full phylogenetic multilevel model including vital-rate CV; thick lines indicate the posterior mean. Panels (a, e) show generation time, (b, f) iteroparity, (c, g) survival unevenness, and (d, h) vital-rate CV. Left y-axes show transient contribution, and right y-axes show log Var(λ), with tick labels shown only on the rightmost panels (d, h).

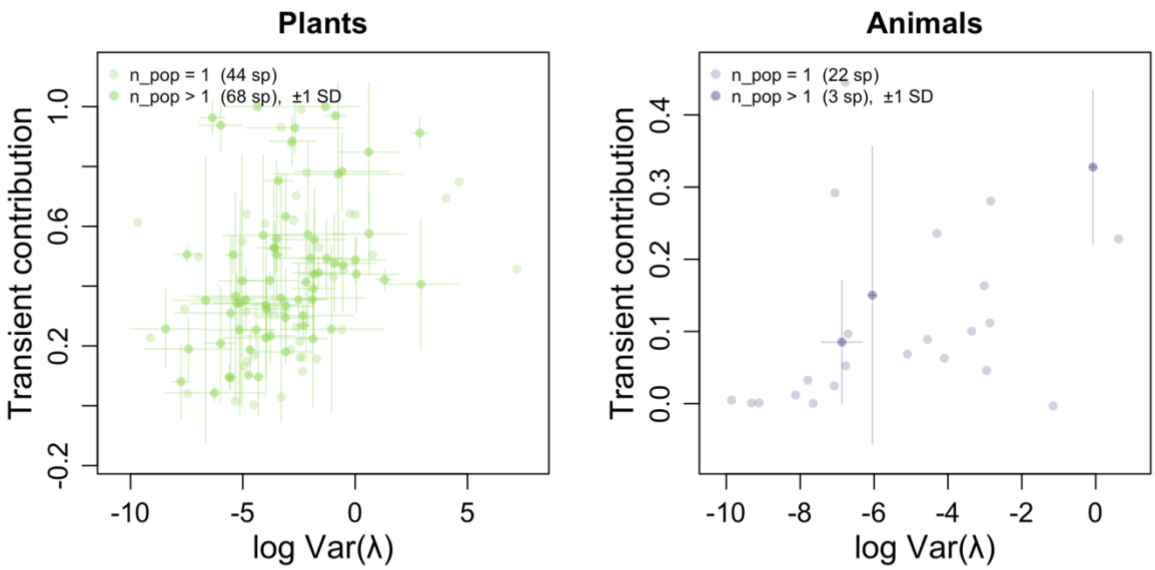


**Supplementary Figure 8. Some plant species show substantial within-species variation in both response variables, while within-species variation is minimal in animals.** Within-species variation in both response variables is present but variable in magnitude across species. Species-level scatter of log-transformed temporal variation in population growth rate (log Var(λ); x-axis) against the proportional contribution of transient dynamics to that variation (transient contribution; y-axis) for plants (left; green) and animals (right; purple). Each point represents one species, plotted at the mean across all available populations. For species represented by more than one population (plants: 68 of 112 species; animals: 3 of 25 species), cross error bars indicate ±1 S.D. across populations in both axes. Species represented by a single population are shown as plain points without error bars.

### Supplementary Methods

#### Body size data

To ensure that differences in demographic variability reflected life history rather than body size, we included adult body mass as a covariate, as evidence indicates that larger-bodied taxa consistently exhibit lower demographic variability (Cohen et al., 2012; Williams et al., 2021). we obtained adult body size from trait databases. For plants, we extracted maximum height data from MOSAIC (Bernard et al., 2023) and the TRY plant trait database (Kattge et al., 2020). When multiple records were available for the same plant species, we chose the maximum standardised value, as done in the comparative literature (e.g. Díaz et al., 2016). For the 53 plant species lacking height data in these databases, we obtained values from floras, species monographs, or herbarium-based accounts. For animals, we used adult body mass values from the MOSAIC database. For the 10 species lacking body mass data in MOSAIC, we obtained values from peer-reviewed publications. For two coral species, we used colony dry mass; for one fish species, we estimated body mass from published length–weight relationships (Froese, 2005). Data sources for all species are provided in Table S1.

#### Quantifying the life-history traits

We computed all three life-history traits from the mean survival–transition matrix (*U*) and the mean fecundity matrix (*F*) of each population, into which the mean projection matrix decomposes as $A = U + F$ (Caswell, 2001). Generation time (*T*) is the mean age difference between parents and their offspring (Bienvenu & Legendre, 2015), obtained from *U* and *F* using the age-difference method (implemented in the R package *Rage*; Jones et al., 2022). Degree of iteroparity (*S*) is Demetrius' entropy (Demetrius, 1978): from *U* and *F* we derived the age-specific survivorship $l_x$ and fecundity $m_x$, defined the share of lifetime reproduction realised at age *x* as $p_x = \frac{l_x m_x}{\sum_x l_x m_x}$, and computed the entropy $S = -\sum_x p_x \ln p_x$ (with the summand taken as 0 when $p_x$ = 0), so that larger *S* indicates reproduction spread more evenly across ages. Survival unevenness ($V_\sigma$) is the coefficient of variation of stage-specific survival

$\sigma_j$ (the column sums of *U*): $V_\sigma = \frac{\mathrm{SD}(\sigma_j)}{\mathrm{mean}(\sigma_j)}$. Higher *Vσ* indicates greater heterogeneity in survival across the life cycle.

**Estimating the variance decomposition using simulations**

To estimate the contributions of the variation in vital rates and population structure to population growth rate, we ran 500 simulation replicates. Specifically, we repeated the simulation procedure across 500 independent replicates for each population and used the resulting distributions, as recommend by Hernández et al. (2026). For each replicate run, we simulated stochastic environmental variation in vital rates using the set of annual transition matrices available for the population (e.g., repeated observations from the same site across multiple years); we began from an initial population vector evenly distributed across classes and applied a burn-in period to avoid any transient effects of the initial population structure. This burn-in period is sufficiently long to remove any effects of the initial population vector (Hernández et al., 2026). We then recorded the vital rates at the final time step and population structure (*i.e.*, population vector) before the final time step to calculate the realised population growth rate ($\lambda_t$). Finally, we calculated the contribution from variation in vital rates to variation in population growth rate ($\mathrm{Var}(\Delta R)$) using the vital rates at the final time step, mean vital rates, and the average stage distribution across simulations. Because the terms in the decomposition of realised population growth rate sum to $\lambda_t$ , the remaining deviation from the asymptotic growth rate (*i.e.*, $\lambda_t - \lambda_d - \Delta R$) corresponds to the contribution from transient dynamics ($V_{\mathrm{transient}}$). This contribution includes both the direct effect of deviations in population structure and their interaction with variation in vital rates. Treating these quantities as random variables across replicate simulations, we estimated their variances and covariances to obtain the variance decomposition described in Methods.

**Phylogenetic corrections**

To account for phylogenetic non-independence among species in our study, we used phylogenies for animals and plants. For plants, we constructed phylogenetic trees using the

*V. PhyloMaker2* package (Jin & Qian, 2022) with the GBOTB extended mega-tree under scenario S3. This scenario inserts unmatched species at the midpoint of their genus or family stem branch where possible. For animals, we obtained a phylogenetic topology using the *rotl* package (Michonneau et al., 2016), which queries the Open Tree of Life synthetic tree and returns an induced subtree for a given set of species. This approach provided substantially broader taxonomic coverage than time-calibrated summary methods, accommodating invertebrate taxa alongside vertebrates. As the Open Tree of Life topology does not carry branch lengths, we imputed branch lengths using the Grafen's transformation (1989), which sets branch lengths proportional to the number of descendent tips at each node. We standardised species names prior to tree construction to reconcile taxonomic synonyms and subspecies designations with accepted names in the reference tree. To incorporate phylogenetic structure into our statistical models, we calculated phylogenetic covariance matrices from the resulting trees using the function *vcv.phylo* from the R package *ape* (Paradis et al., 2004), and incorporated these matrices as a species-level random effect in our multivariate Bayesian models, below.

Before fitting the full models, we assessed phylogenetic signal in each response variable using intercept-only models. Signal was negligible in plants (Pagel's $\lambda = 0.04$ for temporal variation in growth rate and 0.03 for the contribution of transient dynamics) but higher and highly uncertain in animals (0.85 and 0.40, respectively), with posterior distributions spanning much of the parameter range (Fig. S3). We therefore report phylogenetically corrected models in the main text and uncorrected models in the Fig. S4–S5.

To quantify phylogenetic signal in each response variable, we fitted intercept-only versions of each phylogenetic model, retaining the phylogenetic random intercept specified via the *gr()* covariance formulation in *brms* (Bürkner, 2021). We estimated phylogenetic signal as an analogue of Pagel's λ (*i.e.,* the proportion of total variance attributable to phylogenetic covariance), and derived its posterior distribution using the *hypothesis()* function in brms (Hadfield & Nakagawa, 2010).

**Priors, collinearity and sensitivity analyses**

We fitted models using Student-t response distributions to improve robustness to extreme observations (Lange et al., 1989). To regularise parameter estimates while avoiding strong assumptions about effect sizes, we assigned weakly informative priors throughout the analyses (Lemoine, 2019). Specifically, we assigned normal priors centred at zero to regression coefficients and intercepts (McElreath, 2018), exponential priors to variance parameters (Gelman, 2006), and a gamma prior to the Student-t shape parameter (Chung et al., 2013).

Prior to model fitting, we assessed collinearity among predictors using variance inflation factors (VIFs). All VIF values were below 3, except for generation time in animal (VIF = 3.04), indicating generally low collinearity among predictors (Zuur et al., 2010). We additionally evaluated the robustness of trait effects using leave-one-out sensitivity analyses in which we sequentially excluded each focal life history trait from the full model while retaining all covariates. We present the results of these sensitivity analyses in Fig. S6.

### Reference

Bernard, C., Santos, G. S., Deere, J. A., Rodriguez-Caro, R., Capdevila, P., Kusch, E., Gascoigne, S. J., Jackson, J., & Salguero-Gómez, R. (2023). MOSAIC-A unified trait database to complement structured population models. Scientific Data, 10(1), 335.

Bienvenu, F., & Legendre, S. (2015). A new approach to the generation time in matrix population models. The American Naturalist, 185(6), 834–843.

Bürkner, P.-C. (2021). Bayesian item response modeling in R with brms and Stan. Journal of Statistical Software, 100, 1–54.

Caswell, H. (2001). Matrix population models. Sinauer Sunderland.

Chung, Y., Rabe-Hesketh, S., Dorie, V., Gelman, A., & Liu, J. (2013). A nondegenerate penalized likelihood estimator for variance parameters in multilevel models. Psychometrika, 78(4), 685–709.

Cohen, J. E., Xu, M., & Schuster, W. S. (2012). Allometric scaling of population variance with mean body size is predicted from Taylor's law and density-mass allometry. Proceedings of the National Academy of Sciences, 109(39), 15829–15834.

Demetrius, L. (1978). Adaptive value, entropy and survivorship curves. Nature, 275(5677), 213–214.

Díaz, S., Kattge, J., Cornelissen, J. H., Wright, I. J., Lavorel, S., Dray, S., Reu, B., Kleyer, M., Wirth, C., & Colin Prentice, I. (2016). The global spectrum of plant form and function. Nature, 529(7585), 167–171.

Froese, R. (2005). FishBase. World wide web electronic publication. Http://Www. Fishbase. Org.

Gelman, A. (2006). Prior distributions for variance parameters in hierarchical models (comment on article by Browne and Draper). Bayesian Analysis, 1(3), 515–534.

Grafen, A. (1989). The phylogenetic regression. Philosophical Transactions of the Royal Society of London. B, Biological Sciences, 326(1233), 119–157.

Hadfield, J. D., & Nakagawa, S. (2010). General quantitative genetic methods for comparative biology: Phylogenies, taxonomies and multi-trait models for continuous and categorical characters. Journal of Evolutionary Biology, 23(3), 494–508.

Hernández, C. M., Jaggi, H., Cant, J., Zuo, W., Tuljapurkar, S., & Salguero-Gomez, R. (2026). A robust method for quantifying the contribution of transient dynamics to variation in population growth rate.

Jin, Y., & Qian, H. (2022). V. PhyloMaker2: An updated and enlarged R package that can generate very large phylogenies for vascular plants. Plant Diversity, 44(4), 335–339.

Jones, O. R., Barks, P., Stott, I., James, T. D., Levin, S., Petry, W. K., Capdevila, P., Che-Castaldo, J., Jackson, J., Römer, G., Schuette, C., Thomas, C. C., & Salguero-Gómez, R. (2022). Rcompadre and Rage—Two R packages to facilitate the use of the COMPADRE and COMADRE databases and calculation of life-history traits from matrix population models. Methods in Ecology and Evolution, 13(4), 770–781.

Kattge, J., Bönisch, G., Díaz, S., Lavorel, S., Prentice, I. C., Leadley, P., Tautenhahn, S., Werner, G. D., Aakala, T., & Abedi, M. (2020). TRY plant trait database–enhanced coverage and open access. Global Change Biology, 26(1), 119–188.

Lange, K. L., Little, R. J., & Taylor, J. M. (1989). Robust statistical modeling using the t distribution. Journal of the American Statistical Association, 84(408), 881–896.

Lemoine, N. P. (2019). Moving beyond noninformative priors: Why and how to choose weakly informative priors in Bayesian analyses. Oikos, 128(7), 912–928.

McElreath, R. (2018). Statistical rethinking: A Bayesian course with examples in R and Stan. Chapman and Hall/CRC.

Michonneau, F., Brown, J. W., & Winter, D. J. (2016). rotl: An R package to interact with the Open Tree of Life data. Methods in Ecology and Evolution, 7(12), 1476–1481.

Paradis, E., Claude, J., & Strimmer, K. (2004). APE: Analyses of phylogenetics and evolution in R language. Bioinformatics, 20(2), 289–290.

Williams, N. F., McRae, L., Freeman, R., Capdevila, P., & Clements, C. F. (2021). Scaling the extinction vortex: Body size as a predictor of population dynamics close to extinction events. Ecology and Evolution, 11(11), 7069–7079.

Zuur, A. F., Ieno, E. N., & Elphick, C. S. (2010). A protocol for data exploration to avoid common statistical problems. Methods in Ecology and Evolution, 1(1), 3–14.
